# Motor cortical dynamics are shaped by multiple distinct subspaces during naturalistic behavior

**DOI:** 10.1101/2020.07.30.228767

**Authors:** Matthew G. Perich, Sara Conti, Marion Badi, Andrew Bogaard, Beatrice Barra, Kanaka Rajan, Jocelyne Bloch, Gregoire Courtine, Silvestro Micera, Marco Capogrosso, Tomislav Milekovic

**Author notes:** these authors jointly supervised this work (Marco Capogrosso, Tomislav Milekovic).

## Abstract

Behavior relies on continuous influx of sensory information about the body. In primates, motor cortex must integrate somatic feedback to accurately reach and manipulate objects. Yet, prior work demonstrates that motor cortex is well-described with deterministic, rather than input-driven, dynamics. Deterministic dynamics facilitate robust movement generation, but flexible motor output requires rapid responses to unexpected inputs. Here, we resolved this paradox by simultaneously recording neural population activity in motor and somatosensory cortex from four monkeys performing a naturalistic object interaction behavior resulting in occasional errors. Motor cortex was strikingly input-driven surrounding behavioral error correction. Intriguingly, input-driven dynamics were isolated to a subspace of the population activity that putatively captured somatosensory feedback. Using electrical stimulation of ascending somatosensory tracts, we causally verified that this feedback subspace captured peripheral inputs to cortex. Our results demonstrate that cortical activity is compartmentalized within distinct subspaces, enabling flexible integration of salient inputs for robust behavior.

## INTRODUCTION

Behavior relies on continuous influx of sensory information about the body and the environment. In primates, cortex must integrate somatic feedback to accurately reach and manipulate objects^1,2^. The motor cortex is an appealing hub for such integration: it receives constant input from sensory cortical areas as it generates behavior^3^ and its neurons respond rapidly to limb perturbations and other sensory inputs^4,5^. Yet, in many experimental regimes, motor cortex is seemingly well-described as a feedforward system with deterministic dynamics^6,7^, rather than an input-driven^3^ system responding to behaviorally-relevant inputs^3^. While such deterministic dynamics can facilitate robust movement generation^8,9^, the ability to rapidly respond to unexpected inputs is critical to adjust motor output in response to errors. Here, we resolved this paradox by studying scenarios where motor cortical dynamics must change unexpectedly to generate accurate behavioral output^1^. We simultaneously recorded activity of neural populations in motor and somatosensory cortex from four monkeys performing a naturalistic object interaction behavior resulting in occasional errors^10^. Motor cortex typically exhibited robust deterministic dynamics^8,9^, yet was strikingly input-driven surrounding correction of behavioral errors. To determine if the input-driven dynamics could be isolated within the population activity, we decomposed motor cortical activity into orthogonal subspaces^11–13^ putatively capturing feedback from the somatosensory cortex or ongoing dynamics related to behavioral output. Only the feedback subspace expressed input-driven dynamics during error correction, while the behavioral subspace maintained deterministic dynamics. Using electrical stimulation of ascending somatosensory pathways in the spinal cord, we causally verified that only the feedback subspace captures peripheral inputs to cortex. We therefore demonstrate that cortical activity is compartmentalized within distinct subspaces that together shape the population dynamics, enabling flexible integration of salient inputs with ongoing activity for robust behavior.

Movement requires the coordination of large populations of neurons across the motor cortex to produce the necessary patterns of activity through the descending corticospinal tract^14–18^. During reaching and grasping movements, a large part of this motor cortical population activity can be well-described as a dynamical system^6,7^ operating within a neural manifold^12,19–23^, where the future state of the neural population activity evolves deterministically from the present state. Thus, it may seem that motor cortical activity can be appropriately viewed as a deterministic generator of movement – i.e. a system that is not strongly influenced by external inputs^7,9^. However, integration of sensory feedback is essential for movement (**Fig. 1a**). Motor cortical activity in primates is shaped by somatic feedback^1^, and the motor cortex in mice requires constant input as it generates behavior^3^. Given that motor cortex can operate as a feedback controller^1,4^, why do the motor cortical dynamics appear to be so deterministic and not input-driven? Even a system with input-driven dynamics may appear to have deterministic dynamics if its inputs match the predictions of the current state of the population activity^8,24^. We would expect this to be the case during planned, practiced movements without errors or perturbations, such as the types of movements typically used in behavioral neuroscience experiments^6,8,15^. Throughout these movements, inputs such as feedback from the limb largely match what is expected from the outgoing motor commands. Therefore, the motor cortical neural activity exhibits deterministic dynamics even though constantly changing inputs are present^8^. Indeed, a correspondence between the expected and received feedback is essential to learn how to accurately control our limbs^25^. The arrival of unexpected inputs – such as somatic feedback from behavioral errors that deviate from the desired motor output – are necessary to determine the presence of input-driven dynamics in the same cortical population (**Fig. 1b**).

**Figure 1.**
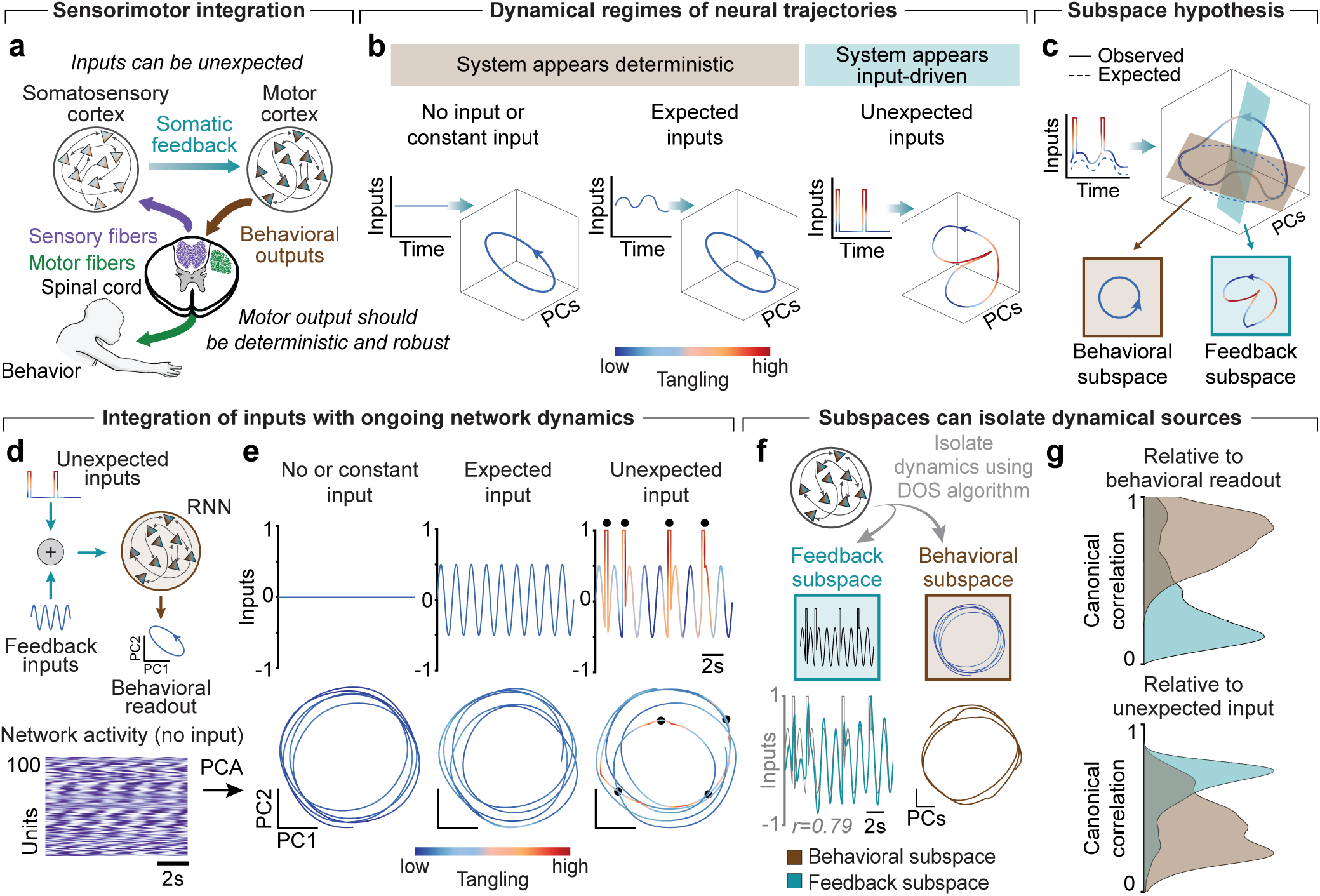
Hypothesis for integrating unexpected inputs with ongoing population activity via subspaces. **(a)** Simplified schematic of interactions between the motor cortex, somatosensory cortex, and behavior. Motor cortex orchestrates the behavior through its behavioral outputs to the spinal cord. Somatosensory cortex receives sensory inputs about the state of the body from the spinal cord. Sensory feedback is communicated to motor cortex from somatosensory cortex to monitor behavioral output. This loop enables generation of robust behavior even in the presence of unexpected inputs, such as caused by behavioral errors. **(b)** Cortical dynamics can appear to be deterministic even in the face of constantly changing inputs so long as those inputs remain predictable or expected. Tangling quantifies the dynamics that a system is currently exhibiting (low tangling: deterministic dynamics; high tangling: input-driven dynamics) and can measure the influence of unexpected inputs on cortical activity. Unexpected inputs can increase tangling in the population dynamics, particularly around the time that the unexpected inputs arrive. **(c)** When all feedback inputs match the expectations (dashed line), motor cortex neural population can exhibit deterministic dynamics. However, when the observed inputs (solid line) deviate from expectations (dashed line), the same population can exhibit input-driven dynamics. We hypothesized that the motor cortex neural population integrates unexpected inputs while maintaining robust movement generation activity through orthogonal subspaces within the neural population activity that respectively exhibit distinct dynamics. **(d)** (Top) We used recurrent neural network (RNN) models to validate the consistency of our hypothesis. We generated 100-unit RNNs with random connectivity and drove them with specific input patterns. We defined the behavioral readout of the RNN as the two leading Principal Components of the full RNN population activity. The input patterns included no input or constant input, as well as input mimicking naturalistic scenarios composed of feedback inputs that reflect the behavioral readout with or without the unexpected inputs. (Bottom) With no input, the network spontaneously generated oscillations, thus resulting in deterministic dynamics. **(e)** With no input or constant input, the behavioral readout produced untangled rotations. With an expected sinusoid input, the behavioral readout remained oscillatory and largely untangled. When driven by a sinusoid that is occasionally unexpectedly interrupted by impulse signals, the behavioral readout was largely untangled except near the unexpected impulse signals marked by the black dots. **(f)** (Top) We then used the RNN model to demonstrate how subspaces can simplify the task of reading out activity related to specific dynamics. We applied our DOS algorithm (see Methods) to isolate behavioral and feedback subspaces. (Bottom) The graphs show projections of neural population activity onto the decomposed input-driven (left) and deterministic (right) subspaces for a single simulation. **(g)** We ran 1000 simulations of RNNs with random connectivity driven by the inputs shown in Panel e. In each simulation, we computed the canonical correlation of the behavioral subspace (brown) and feedback subspace (blue) with the RNN’s behavioral readout (Top) and the unexpected input (Bottom). ***: p < 0.001, rank-sum test.

Prior analyses of motor cortical dynamics have used repetitive or highly trained behavior and found it to be surprisingly deterministic^6–9^. The deterministic dynamics can be advantageous for the motor cortex as it allows generation of robust control signals that are resistant to noise^9^, including the biological limitations of neural circuits^8^ such as neuronal death^26^ or unreliable synapses^27^. Can the motor cortex integrate relevant inputs while maintaining deterministic dynamics to robustly generate behavior, or is the population of neurons forced to switch between deterministic and input-driven regimes? We hypothesized that these deterministic and input-driven dynamics can effectively co-exist if they are confined to independent dimensions within the neural manifold otherwise known as *subspaces*. This framework allows cortex to operate as a set of linked but independently-driven systems, one comprising input-driven and another deterministic dynamics. Together, these systems shape the global cortical dynamics observed in the full population (**Fig. 1c**).

### Simulations reveal that a neural network can reproduce multiple dynamical regimes

To motivate our hypothesis, we designed a simulation in which a recurrent neural network (RNN) with random connectivity is driven by external inputs. We defined a behavioral readout of the RNN to be the two leading principal components of the full RNN population activity. The inputs could be considered either expected or unexpected based on the oscillatory patterns in the current RNN behavioral readout. We used the metric of *trajectory tangling*^9^ to quantify the degree to which the network momentarily exhibits deterministic or input-driven dynamics. Tangling is low when similar neural states – here, the position and velocity of the neural trajectory in state space – evolve similarly and increases when evolution of similar neural states diverge. For our purposes, we interpret periods of high tangling to indicate that a system is transiently driven by unexpected inputs that cause the trajectory to deviate from the predicted, deterministic path. In contrast, we interpret low tangling to indicate a system in which current activity uniquely predicts future activity, i.e. a system exhibiting deterministic dynamics. Note that low tangling need not imply that no inputs are driving the system. As described above, if the inputs match the expectations of the outgoing motor command, then the evolution of neural states will be quantified by low tangling.

Thus, even a system which is always input-driven can sometimes exhibit low tangling and unveil higher tangling when the inputs differ from the expectation (**Fig. 1b**).

In the absence of feedback inputs, the RNN activity produced oscillations with low tangling, thereby indicating deterministic dynamics (**Fig. 1d**). When we introduced an expected sinusoid input, the RNN trajectory still evolved in a deterministic manner and its tangling remained similarly low. If the activity of RNN units represented recordings taken from a brain region during behavior, analysis of this trajectory alone would not be sufficient to indicate whether the population exhibited strictly deterministic dynamics, input-driven dynamics, or incorporated both dynamical regimes. Separating these possibilities requires new inputs that will cause neural states to diverge from expected paths. When we simulated such inputs to the RNN by adding transient impulses to the sinusoid input, the tangling of the RNN trajectory drastically increased around the times of the impulses (**Fig. 1e**). This result demonstrated that RNNs can indeed reproduce both dynamical regimes depending on the type of inputs. Furthermore, this simulation underscores the importance of providing rich behavioral contexts with varied, unexpected inputs (i.e., system perturbations) to identify the presence of deterministic and input-driven dynamics in a neural population. Nonetheless, it remained unclear whether RNNs have to switch between deterministic and input-driven dynamics, or if deterministic and input-driven dynamics are separated into linearly separable subspaces.

### Decomposing population activity into subspaces

As outlined before, the generation of behavior can benefit from robust, deterministic dynamics. Yet, input-driven dynamics are necessary to receive feedback about ongoing movements. When performing a motor task in a natural environment, this input is expected to arrive from or at least coincide with activity of the primary somatosensory cortex. Recent work has demonstrated that the neural manifold can be decomposed into *subspaces* capturing specific population-wide features. Subspaces can capture planning a movement^12,19^, adjusting motor output during learning^11,28^, sharing the control of a limb across hemispheres^29,30^, and even communicating between cortical regions^13^. Here, we hypothesize that linear subspaces, hereby named *feedback* and *behavioral* subspaces, can sufficiently separate independent dimensions containing input-driven and deterministic dynamics related to somatosensory feedback and behavior, respectively. This hypothesis can be directly validated experimentally – if true, during a motor task all unexpected somatosensory inputs will predominantly influence the feedback subspace activity, thus increasing its tangling, while the behavioral subspace tangling will remain unchanged. We predicted that motor cortical activity that covaries with either the behavior or the activity of the somatosensory cortex can aid in the identification of the motor cortex behavioral and feedback subspaces, respectively.

We provided proof of principle of this prediction using our RNN model. We developed an algorithm – Decomposition into Optimal Subspaces (DOS) – to identify these subspaces within the population activity (see Methods). This algorithm optimally decomposes neural population activity into orthogonal subspaces based on the covariance of the neural population activity with other variables, such as behavior or the activity of another neuronal population. We first verified that DOS accurately isolates specific dynamics from simulated datasets with known ground truth dynamics (**Fig. S1;** see Methods). We then tested whether the input-driven and deterministic components of the RNN trajectories can be linearly separated into subspaces (**Fig. 1f**). We found that a linear decomposition sufficed to isolate dynamics that were highly correlated with the empirical deterministic dynamics (**Fig. 1g**), in a manner that was isolated from the input-driven dynamics. This simulation illustrates the utility of subspaces for isolating and understanding the dynamical regimes describing a neural trajectory. We next applied these concepts to cortical recordings during a naturalistic behavior to determine how deterministic and input-driven dynamics can co-exist within cortical populations.

### A naturalistic reach, grasp and manipulate task to study sensorimotor integration

The surprisingly deterministic nature of motor cortical dynamics in prior experiments may be attributed to studying highly-trained subjects performing planned or repetitive movements^6,7,22^. To understand the effect of unexpected feedback on motor cortical dynamics, we designed a naturalistic task that resulted with occasional behavioral errors. In our task, monkeys reached for, grasped, and pulled objects mounted on a robotic arm^10^ (**Fig. 2a**), producing a naturalistic behavior similar to picking fruit from a tree. During the pulling phase, the robot acted as a spring, applying a resistive force proportional to the displacement. This task had several unique properties: 1) the initial reaching phase was similar to many prior studies^6,12,31^; 2) object contact (i.e. grasp) provided a salient sensory cue; and 3) the resistive pulling phase required manipulation of an external object.

**Figure 2.**
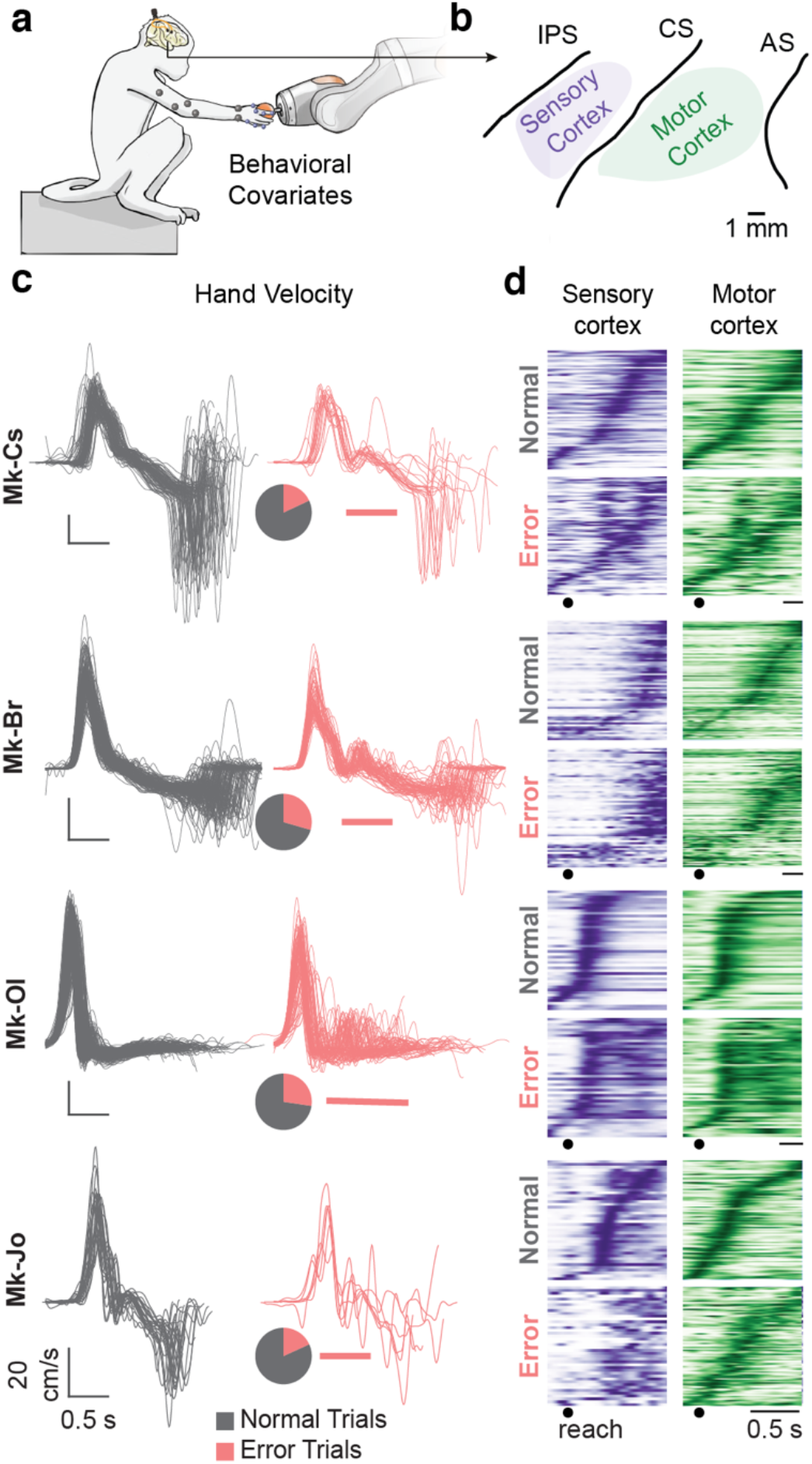
Behavioral task and recordings. **(a)** Monkeys reached for, grasped, and pulled an object mounted on a haptic robot while we recorded limb kinematics, pulling forces, and neural population activity from motor and somatosensory cortex simultaneously. **(b)** Approximate recording locations for all four monkeys spanned between the central sulcus (CS) and intraparietal sulcus (IPS) for somatosensory cortex (purple), and the CS and arcuate sulcus (AS) for motor cortex (green). **(c)** Plots show the hand velocity in the sagittal plane for all normal (gray, left) and error (pink, right) trials for each of the four monkeys. Pie charts show the proportion of error trials: 18%, 29%, 27%, and 18% of trials for Mk-Cs, Mk-Br, Mk-Ol, and Mk-Jo, respectively. Each behavioral error required a behavioral correction characterized by a second period of forward velocity to either reach for the robot again or adjust the position of the hand. Pink bars beneath the trajectories highlight the approximate window of error correction for each monkey. **(d)** Average neural activity for all somatosensory (purple) and motor (green) cortical neurons for each monkey for normal (top) and error (bottom) trials. We isolated 78, 126, 62, and 88 motor and 82, 59, 44, and 42 somatosensory cortical neurons from Mk-Cs, Mk-Br, Mk-Ol, and Mk-Jo, respectively. Activity was aligned on reach and averaged across trials. Neurons are sorted based on their time of maximal activity in normal trials.

We trained four monkeys (*macaca fascicularis*) to complete this task. Each monkey exhibited an idiosyncratic behavioral strategy (**Fig. 2c and S2**). However, all monkeys occasionally made behavioral errors – that is, deviations from an optimal reach, grasp, and pull strategy comprising smooth continuous movements – before ultimately succeeding in pulling the object. For Mk-Br, most errors occurred after grasping the object to adjust the posture of the hand to enable the pulling movement. For the remaining monkeys, errors were typically due to the hand missing the object or slipping off the object as it was pulled. These monkeys compensated by adjusting the position of the hand and grasping and pulling again (**Fig. 2c**). These error trials provided an opportunity to study the effect of unexpected inputs on cortical activity.

As the monkeys performed the task, we simultaneously recorded kinematics of the left arm (Vicon, Oxford, UK), three-dimensional pulling force on the object, and the spiking activity of motor and somatosensory cortical neural populations (**Fig. 2b**) using chronically-implanted electrode arrays (Blackrock Microsystems, Salt Lake City, UT, USA). When comparing the normal to error trials, we saw a transient reorganization of activity in both motor and somatosensory cortex around the time of error correction (**Fig. 2d**). For example, Mk-Cs (**Fig. 2d**) had a prominent burst in activity in both regions around the time of error correction and subsequent pulling phase. These bursts likely resulted from unexpected inputs arriving to both motor and somatosensory cortex reflecting the occurrence of the error, as well as altered activity driving the behavioral correction.

### Motor and somatosensory cortex exhibit distinct dynamics during reach to grasp

To characterize the dynamics of the somatosensory and motor cortical population activity, we applied Principal Component Analysis (PCA) to identify 20-dimensional manifolds^20^ that captured between 68-86% of neural population variance in each cortical region. We also applied PCA on the recorded arm kinematics and pulling force (behavioral covariates) to identify a 20-dimensional “behavioral manifold” that captured at least 95% of the variance. To quantify the local dynamical regime of each manifold, we applied the trajectory tangling metric described above^9^. However, tangling is a relative metric and can be challenging to concretely interpret in the context of input-driven or deterministic systems. To set a reference for tangling values, we used RNN models directly constrained by the neural population recordings that recapitulated the joint motor and somatosensory cortical recordings as a fully autonomous (i.e., not input-driven), deterministic dynamical systems using only the initial condition on each trial^32^ (**Fig. S3**). Since the RNNs were autonomous by construction, they were only capable of capturing the portion of the population dynamics that can be readily explained as deterministic. By computing the distribution of tangling values from these model RNNs, we were able to identify a range of tangling values for each individual monkey’s dataset that would be consistent with purely deterministic dynamics (colorbars in **Fig. 3e**).

**Figure 3.**
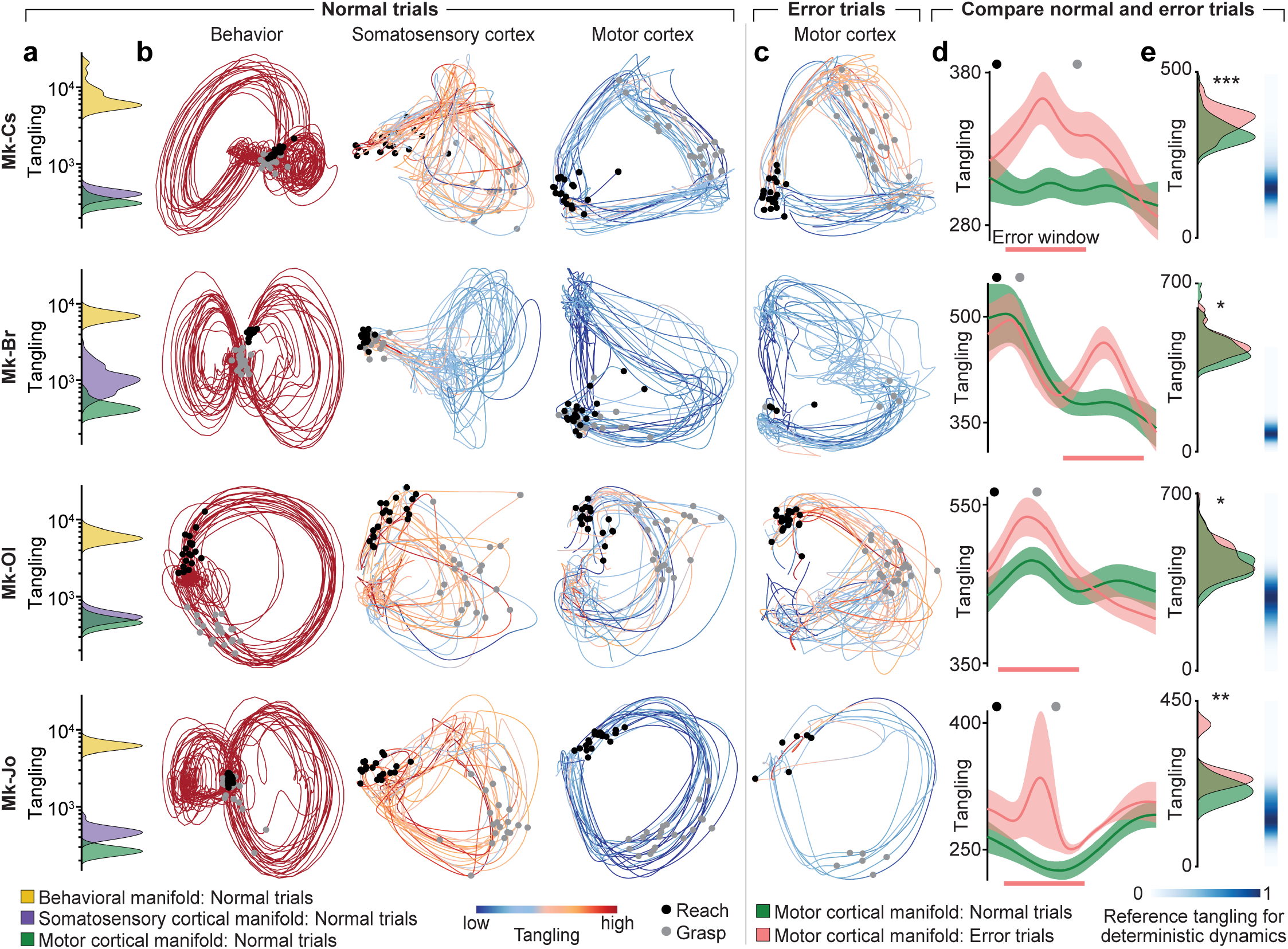
Motor cortical population activity is driven by unexpected inputs on error trials. **(a)** Tangling of the behavior manifold is substantially higher than the tangling of the somatosensory cortical manifold, which is higher than the tangling of the motor cortical manifold. We identified 20 dimensional manifolds capturing the behavioral signals (yellow), as well as the somatosensory (purple) and motor (green) cortical population activity. We then computed the tangling of the trajectories of behavior, somatosensory cortical and motor cortical manifolds. The distributions show mean tangling across a trial for all trials in each of these three manifolds on a log scale. All distributions are significantly different at p < 0.001 (ranksum). **(b)** Each subplot shows the trajectories in the two leading Principal Components of the behavioral (left), somatosensory cortical (middle), and motor cortical (right) manifolds for 20 normal trials. Colors indicate tangling value at each time point on a logarithmic scale, where blue corresponds to the minimum and red corresponds to the maximum neural tangling across all time points of all normal and error trials for each monkey (allowing behavior manifold values to saturate). Black circles: reach onset; gray circles: object grasp. **(c)** Tangling in the motor cortical manifold increases in the error trials. Plots show motor cortical manifold trajectories and tangling for all error trials for each of the four monkeys. Color scale for each monkey is the same as in Panel b. Note that the states at the time of object grasp are similar on the normal and error trials, indicating that error correction required a return to a similar neural state to complete the trial. **(d)** Single-trial error-related motor cortical manifold tangling is significantly higher in error trials compared to the normal trials. Tangling is computed in windows shown by pink lines in Panel e. Plots show the distributions across single trials of average tangling in the motor cortical manifold for all normal (green) and error (pink) trials. Black and grey circles indicate the mean time of reach onset and object grasp, respectively. *: p < 0.05, **: p < 0.01, ***: p < 0.001; ranksum. **(e)** Motor cortical manifold tangling acutely increases surrounding the error correction. Plots show mean of tangling in the motor cortical manifold across all normal or error trials after aligning on reach onset and time-warping to match the length of each reach and pull phase across all trials (see Methods). Pink bars indicate approximate windows of error correction for each monkey. We used RNN models directly constrained by the motor cortex manifold activity as a reference system exhibiting exclusively deterministic dynamics. We computed the distribution of tangling values from 1000 of these randomly-initialized model RNNs to map tangling values specific to that motor cortex manifold that are consistent with purely deterministic dynamics. Blue color gradients show the distribution of these tangling values in the scale of the right column of panels. Error bars: mean ± s.e.m.

The trajectories in the behavioral manifold had consistently very high tangling for all four monkeys (**Fig. 3a,b**). These trajectories typically exhibited a cross-over point at reach onset and grasp, where most of behavioral variables are similar (hand at rest). Although somatosensory cortical activity is largely driven by feedback about the behavioral output, the trajectories of the somatosensory manifold (**Fig. 3a,b**) were substantially less tangled than in the behavioral manifold. The cross-over point was absent, yet the trajectories made sharp turns at different points, thereby causing the paths to frequently diverge and converge resulting in increased tangling. Despite the apparent similarities in the firing patterns of somatosensory and motor cortical neurons (**Fig. 2d and S2d-e**), trajectories in the motor cortical manifold were circular and smooth (**Fig. 3a,b**) resulting in significantly lower tangling than in both behavioral and somatosensory cortical manifolds^9^. When we derived estimates of purely deterministic dynamics from the data driven RNN models (**Fig. S3**), we found that motor cortical manifold tangling values were uniformly these values (**Fig. 3e**, blue gradients) indicating that it is likely operating in an input-driven regime.

We then analyzed the manifold trajectories on error trials. We predicted that tangling of the motor cortex manifold trajectory should increase when sensory inputs indicate a need to alter the current behavior, such as following the behavioral errors. In our task, each error was followed by a behavioral correction to complete the trial (**Fig. 2b**). These corrections corresponded to loops or distortions in the motor cortical manifold trajectories (**Fig. 3c**). We saw that motor cortical manifold tangling increased transiently surrounding the error correction (**Fig. 3c-d**), indicating that the motor cortex may then be driven by unexpected inputs. When compared to the reference values for deterministic dynamics derived from using the RNN model (**Fig. S3**), the tangling values around the behavioral errors were well above the region with the predominantly deterministic dynamics (**Fig. 3e**). We observed similar increases in tangling for the somatosensory cortical manifold trajectories following behavioral errors (**Fig. S4, S5**).

### Transient shifts of the motor cortex towards input-driven dynamics during error corrections are isolated to its feedback subspace

Our simulations using RNN models showed that a neural network can compartmentalize deterministic and input-driven dynamics in different subspaces (**Fig. 1f**). This compartmentalization could be useful for the motor cortex: a subspace with deterministic dynamics can help generate robust behavior, while a subspace with input-driven dynamics can readily incorporate inputs from somatosensory cortex, which are critical for dexterous behavior. Therefore, we hypothesized that applying the DOS algorithm to find the motor cortical activity subspaces that covary with the behavior and, respectively, the somatosensory cortex activity will exactly derive the motor cortex behavioral and feedback subspaces. We predicted that the transient increases of tangling observed in the motor cortical activity during the errors (**Fig. 3e**) are confined to the feedback subspace, while the tangling of the behavioral subspace remains unchanged.

To validate our hypothesis, we applied the DOS algorithm on our experimental datasets to identify feedback and behavioral subspaces of the motor cortical manifolds using covariance with the somatosensory cortical manifolds and the behavioral manifolds, respectively (**Fig. 4a and S6a**). Using all normal trials, we computed the neural variance explained by the feedback and behavioral subspaces for assumed dimensionalities between one and five (**Fig. 4b**). For all four monkeys, the feedback subspace explained the majority of neural population variance at all dimensionalities, even though DOS seeks only to explain covariance with somatosensory cortex, not variance in the motor cortical population. Thus, cortico-cortical interactions appear to be a dominant feature of motor cortical population activity during movement. This observation is particularly surprising given the strikingly different dynamics of the full motor and somatosensory cortical manifolds (**Fig. 3a-b**).

**Figure 4.**
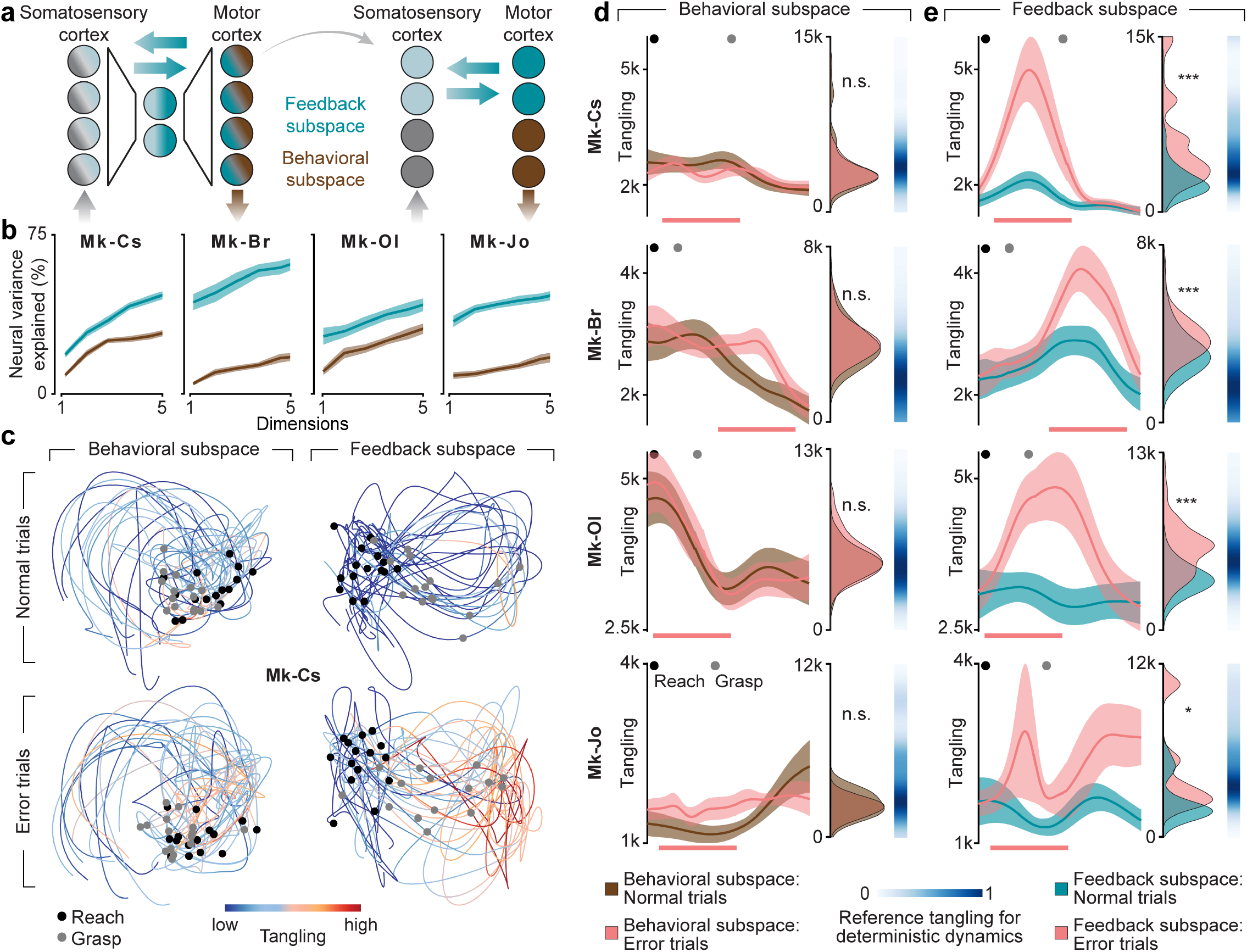
The increase in motor cortical tangling during behavioral errors is confined to its feedback subspace. **(a)** Schematic of the subspace decomposition. The neural manifold identified by PCA comprises a mixture of activity related to cortico-cortical feedback inputs (blue) and behavioral output (brown). We applied the DOS algorithm to identify putative feedback and behavioral subspaces that isolate motor cortex activity related to somatosensory cortical activity or behavioral output, respectively. **(b)** Interactions with somatosensory cortex capture the dominant component of motor cortical activity. The plots show the percentage of motor cortical neural variance explained by the feedback and behavioral subspaces as a function of the chosen subspace dimensionality. **(c)** Feedback subspace tangling substantially increases in error trials. Plots show trajectories within the behavioral (left) and feedback (right) subspaces on normal (top) and error (bottom) trials for Mk-Cs. Data plotted as in Figure 3a. **(d)** Behavioral subspace tangling remains unchanged between normal and error trials. Panels show tangling in the behavioral subspace for all normal (brown) and error (pink) trials for the four monkeys. Left column data are plotted as in Figure 3d. Right column data are plotted as in Figure 3e. We used RNN models to map tangling values specific to that behavioral subspace that are consistent with purely deterministic dynamics (see Fig. 3 for details). Blue color gradients show the distribution of these tangling values in the scale of the right column of panels. **(e)** Feedback subspace tangling substantially increases in the error trials surrounding the error correction. Panels show tangling in the feedback subspace for all normal (blue) and error (pink) trials for the four monkeys. Pink bars show approximate window of error correction. Data presented as in Panel d. Blue color gradients same as in d but for feedback subspace. Error bars: s.e.m. *: p < 0.05, **: p < 0.01, ***: p < 0.001; ranksum.

We next analyzed the dynamics within the motor cortical subspaces using tangling. Since tangling is sensitive to dimensionality of the subspace used to compute the metric, we fixed the dimensionality of the feedback and behavioral subspaces to five to enable direct comparison. Note that tangling by design increases as the dimensionality of the embedding space decreases (**Fig. S7**). Therefore, the tangling values can only be compared across spaces of the same dimensionality. Even on normal trials, the tangling values have a higher baseline value in the five-dimensional subspaces compared to the full twenty-dimensional manifold (**Fig. S7**). Yet, we should still observe significant increases in tangling on the error trials around the times of the error correction behavior even though the dimensionality is lower. If motor cortex receives unexpected inputs from somatosensory cortex during error trials, we hypothesized that the increase in tangling observed in motor cortical activity is isolated to its feedback subspace. To ensure our results were not biased, we projected the activity in error trials into the subspaces previously identified using only the normal trials (**Fig. 4c, Fig. S6**). Despite the increase in tangling observed within the full motor cortical population on error trials (**Fig. 3c-e**), we observed no significant difference in tangling within the behavioral subspace (**Fig. 4d**). Overall, the tangling values within the behavioral subspace were generally within the ranges expected by purely deterministic dynamics derived from our RNN models (**Fig. S3**).

In contrast to the behavioral subspace, trajectories within the feedback subspace increased in tangling around the time of error correction (**Fig. 4e**), similarly to the full motor cortical population activity (**Fig. 3c-e**). This increase in tangling was not observed in the remaining dimensions of population activity that were not captured by either the feedback or behavioral subspaces (the *null subspace*; see Methods), indicating that unexpected somatosensory inputs related to the behavioral errors impact only the feedback subspace dynamics (**Fig. S8**). Between the normal and errors trials, the feedback subspace tangling became substantially higher than the reference deterministic dynamics values obtained using the RNN models of purely deterministic dynamics (**Fig. 4e and S3**). These results together demonstrate that the increased tangling in motor cortex on error trials, indicative of an input-driven regime, is confined to the feedback subspace to allow continually deterministic dynamics to exist within the behavioral subspace. Motor cortical activity is thus composed of multiple, independently-driven systems that together comprise the total dynamics of the region.

### The motor cortical feedback subspace captures direct somatic inputs from ascending spinal sensory tracts

The feedback subspace, as defined in our analysis, captures shared variance between motor and somatosensory cortex, yet this does not on its own show that it necessarily represents somatosensory information from the periphery. To establish this link, we elicited afferent volleys along the ascending dorsal column tracts of the spinal cord by applying epidural electrical stimulation (EES) targeting the dorsal roots innervating cervical segments^33–35^ in two monkeys, Mk-Br and Mk-Yg (**Fig. 5a,b**). Both monkeys received lesions of the descending pathways between C4 and C5 spinal segments (**Fig. 5c**). These lesions spared the voluntary control of proximal muscles that control shoulder and elbow movements but resulted in the paralysis of the distal muscles that control the wrist and finger movements. The electrodes that delivered EES were placed caudal to the lesion to target the C5-T1 dorsal roots. We simultaneously recorded from both motor and somatosensory cortices contralateral to the lesion.

**Figure 5.**
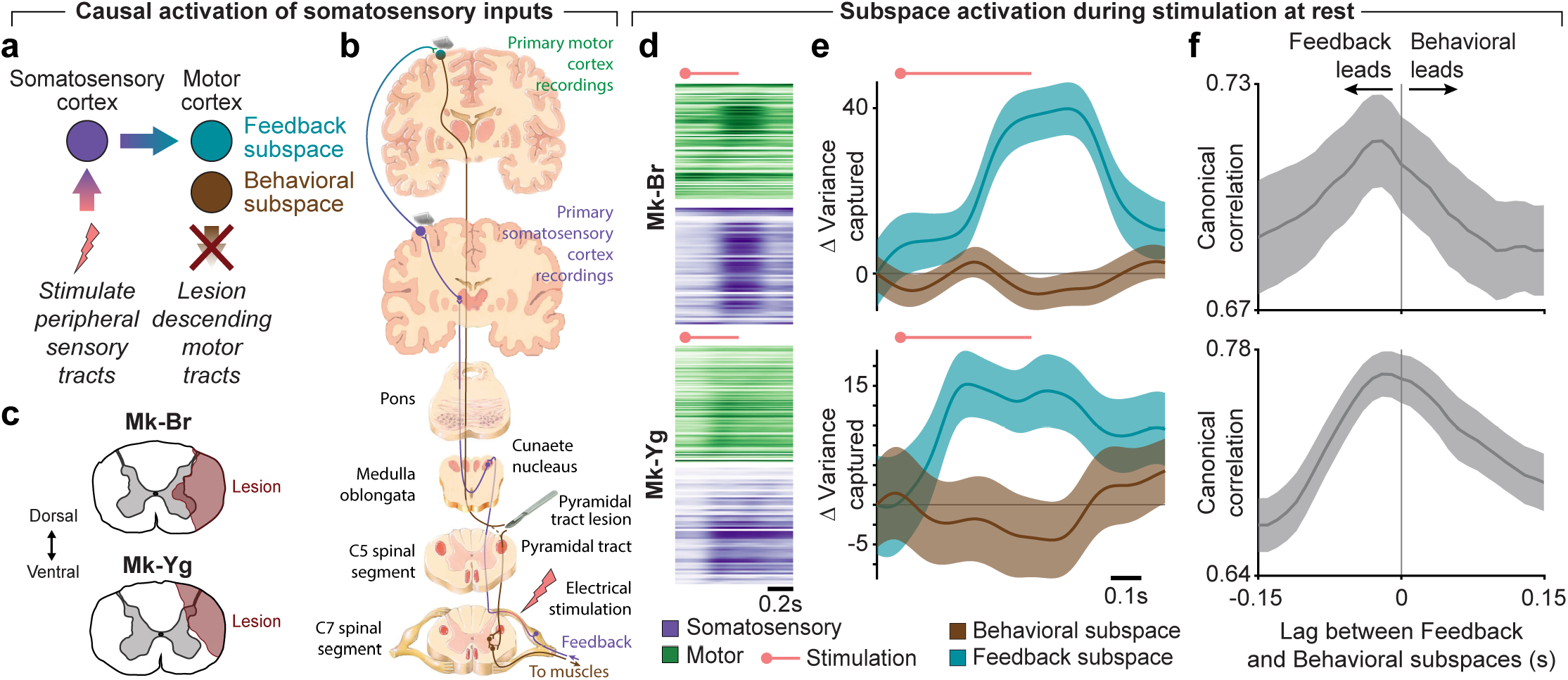
The increase in motor cortical tangling caused by the stimulation of the somatosensory spinal tracts is confined to its feedback subspace. **(a)** We set out to determine how the two motor cortical subspaces respond to the controlled peripheral input. To this end, we stimulated spinal sensorimotor tracts of two monkeys to causally activate cortical circuits through somatic pathways. To ensure activation of only the somatosensory pathways, we lesioned the descending corticospinal tracts above the stimulation site. The stimulation was ipsilateral to and below the lesion. **(b)** We stimulated the dorsal aspect of the spinal cord using flexible electrode arrays inserted under the vertebrae over the C5-T1 spinal segments. We performed a unilateral lesion of the descending cortico-spinal tracts between C4 and C5 spinal segments (red) above the electrode array, severing the majority of the descending motor fibers (green) and preserving the ascending sensory fibers (purple). The lesion was made contralateral to the recorded motor and somatosensory cortices and the stimulation was delivered ipsilateral to the lesion. **(c)** Cross-sectional reconstruction of lesioned areas in both monkeys. **(d)** Average activity of each motor and somatosensory cortical neuron aligned on start of the EES train targeting dorsal roots innervating hand muscles while the monkey was at rest. Both regions responded strongly to the stimulation. The red dot above indicates the onset of stimulation; the line indicates the pulse train duration. **(e)** Change in variance captured after the EES as a function of time relative to the variance captured during rest for the motor cortical behavioral (brown) and feedback (blue) subspaces. **(f)** We computed the canonical correlation between the activity in the behavioral and feedback subspaces at different relative lags in the one second window following stimulation onset. We found the highest correlation in both monkeys when the feedback subspace activity was made to lead the behavioral subspace activity by approximately 30ms.

We first applied trains of EES pulses targeting dorsal roots below the lesion that innervate hand muscles while the monkeys were at rest. These trains elicited action potentials in the proprioceptive axons whose branches join the ascending dorsal column tract^33,34,36^. Activity in the ascending dorsal column tract excites somatosensory cortex neurons via a first relay in the dorsal column nuclei and then a second relay in the thalamus. In turn, the somatosensory cortical neurons directly project to motor cortical neurons^37,38^. As a result, we observed pronounced and sustained responses to EES in both motor and somatosensory cortices (**Fig. 5d**).

We then collected neural and behavioral recordings while the monkeys with impaired wrist and finger movements attempted the object reach, grasp and manipulation task without EES. Since the voluntary control of proximal muscles was spared, the monkeys generated ample movements, which we then used to identify feedback and behavioral subspaces of the motor cortex using the DOS algorithm. We then computed the change in variance of the feedback and behavioral subspaces during EES compared to no stimulation (**Fig. 5e**). We saw a pronounced increase in variance captured by the feedback subspace during EES, with no corresponding increase in the behavioral subspace. This result establishes a causal link between peripheral inputs and the preferential modulation of the feedback subspace. Note that the subspaces were defined during arm movements performed in the complete absence of EES. Consequently, EES served as an independent validation of the subspaces. We then analyzed the temporal relation between the feedback and behavioral subspace responses to EES by computing the canonical correlation between the subspace responses at different temporal shifts (**Fig. 5f**). We found that the responses were most similar when the feedback subspace led the behavioral subspace by approximately 30ms, again consistent with a direct modulation of the feedback subspace by somatosensory inputs from the periphery.

## DISCUSSION

Although inputs, whether sensory^4,39^ or cognitive such as during changes of mind^40,41^, influence motor cortical activity, the mechanisms that allow these inputs to coexist with ongoing population activity driving behavior have remained unclear. Intriguingly, motor cortex is dominated by apparently deterministic dynamics during the execution of planned or repetitive movements^6,8,9^. Modeling provides strong evidence that deterministic dynamics enables motor cortical populations to robustly generate motor output despite biological limitations of the neurons, such as neuronal noise, unreliable synapses or spontaneous death of neurons^8^. Nonetheless, motor cortex has to respond to external inputs when those inputs require unplanned changes of behavior, such as after errors whether self-induced^42^, due to perturbations^28,43^, or during learning^11,31,44,45^. Here, we show how this apparent contradiction may be resolved by compartmentalizing dynamics into lower-dimensional subspaces of the neural manifold: input-driven dynamics within a feedback subspace related to the inputs coming from the somatosensory cortex, and deterministic dynamics within a behavioral subspace related to the motor output.

Although we assume linear subspaces in our analysis, this need not be the case. Indeed, we predict that the linear subspaces we identified are likely an approximation of a true nonlinear manifold^20^. However, the success of our linear decomposition provides an important proof of principle regarding how different dynamical sources can mix in a single neural population, even if the subspaces do not necessarily capture the precise geometric or mechanistic implementation. An interesting open question is how the independently-driven subspaces are linked to allow flexible sensorimotor integration and generate the full, population-wide dynamics. One intriguing possibility is that the link between the subspaces is direct but transient. Another possibility is that the subspaces are perpetually independent, but communication between them is mediated using additional dimensions within the neural manifold^46^ or even through other brain areas. Future theoretical and experimental work is needed to elucidate possible mechanisms underlying this integration.

In this work, we applied the trajectory tangling metric^9^ to distinguish between regimes of input-driven and deterministic dynamics. In our interpretation, this metric provides a convenient, quantitative means to compare differences in neural trajectories. However, tangling is best viewed as a tool for observing the relative dynamics currently exhibited by a system, not an absolute indicator of any one dynamical regime. On error trials, we found that motor cortex exhibited more strongly input-driven dynamics (quantified by an increase in tangling) transiently surrounding the error correction. Surprisingly, this increase in tangling was isolated to the motor cortical feedback subspace. This result cannot be trivially explained. First, we derived the subspaces using only normal trials, ensuring that there was no bias encouraging the feedback subspace to capture the input-driven dynamics on error trials. Second, the tangling measure was not considered when defining the subspaces, ensuring that tangling remains an unbiased metric quantifying the dynamics. Third, the increased tangling is not a trivial consequence of covarying motor cortex with a more tangled cortical region like somatosensory cortex. There was no apparent bias for the feedback subspace to have higher tangling on normal trials, and the covariates used to define the behavioral subspace had the highest tangling overall (**Fig. 3a**). Despite anatomical connections between the regions, our experiments cannot fully implicate somatosensory cortex in driving these new motor cortical dynamics. Indeed, we would expect similar results if both cortical regions were driven simultaneously by a third region such as the thalamus^3^. However, regardless of the exact pathway leading to motor and somatosensory cortex, our spinal stimulation experiment provides evidence that our feedback subspace preferentially captures direct, feedback-related peripheral signals. Experimental tools that facilitate precise and real-time manipulation of specific neural circuit pathways could help in future work to causally demonstrate the role of subspaces for sensorimotor integration.

The activity of many cortical regions can be accurately represented by low-dimensional dynamics, despite the large number of neurons involved in their function^11,20,47–50^. Prior work has shown that untangled population activity allows for robust control signals within motor cortex^9^, but is not necessarily an ubiquitous phenomenon across all cortical regions. Here, we have shown in four monkeys that sensory cortical areas are more tangled than motor areas, as has been previously reported^9^. Yet, our subspace analysis shows that even within a single cortical region the neural population can be decomposed into independently-driven subspaces operating within distinct dynamical regimes that together shape the full population dynamics. Here, we studied the interactions between motor and somatosensory cortex, though the motor cortex also receives input from many other regions in the brain. The high possible dimensionality of cortical populations could enable inputs from many brain regions to be simultaneously integrated with ongoing computations by compartmentalizing them into distinct subspaces. Further experiments that simultaneously record from several interconnected cortical regions are needed to confirm whether cortex utilizes this approach to integrate multiple inter-regional inputs.

## ACKNOWLEDGEMENTS

We thank Maude Delacombaz and Melanie Kaeser for assistance with behavioral training of the animals. M.G.P. thanks Dr. Juan A. Gallego and Dr. Raeed H. Chowdhury for their valuable discussions in the preparation of this manuscript. Funding: Whitaker International Scholars Program fellowship (M.G.P.), Fonds de recherche du Québec Santé (chercheurs-boursiers en intelligence artificielle; M.G.P.), Wyss Center for Bio and Neuroengineering (WCP-008, G.C., M.C. and T.M), ONWARD Medical (G.C, M.C), Bertarelli Foundation Catalyst Fund (BC1709, T.M. and M.C.), the Bertarelli Foundation, and the Swiss National Science Foundation through Ambizione awards (PZ00P2_168103, T.M.; PZ00P2_167912, M.C.), the NeuGrasp project (170032, S.M.), and the Swiss National Centre of Competence in Research (NCCR) Robotics.

## AUTHOR CONTRIBUTIONS

*Conception and theory:* M.G.P.

*Experimental Design:* M.G.P., S.C., M.B., A.B., B.B., J.B., G.C., S.M., M.C., and T.M.

*Surgical implantation:* M.G.P., A.B., J.B., and G.C.

*Animal care:* M.G.P., S.C., M.B., A.B., S.W., and M.C.

*Data collection and processing:* M.G.P., B.B., S.C., M.B., A.B., and M.C.

*Neural data analysis:* M.G.P.

*Design of recurrent neural network models:* M.G.P. and K.R.

*Figure preparation:* M.G.P. and T.M.

*Manuscript preparation:* M.G.P.

*Manuscript and figure editing:* M.G.P., G.C., M.C., and T.M.

## METHODS

### Simulation to study the effect of inputs on tangling

We developed a recurrent neural network (RNN) simulation to study the integration of inputs with ongoing deterministic dynamics. We generated a 100-unit RNN model governed by random connectivity drawn from a Gaussian distribution and scaled by a factor *g* > 1 to induce chaotic oscillations in the network. We computed the change in activity at each time step according to:

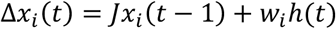

where *J* is the connectivity matrix governing the network which is scaled by *g, xi(t)* is the activation of each unit *i, w* is the input weight to each neuron and *h(t)* is the time-varying input driving the network. The state update of the network was then governed by:

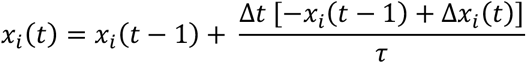

where τ is the decay time constant of the units in the network. In these simulations, we set *g*=1.5 and τ=0.25. We selected various one-dimensional signals with equal connections to all units in the network. We defined the condition where the input was zero to be the “deterministic dynamics”, where the network deterministically generated chaotic oscillations based on the initial condition. We then provided a sinusoid input to the network to demonstrate the effect of expected time-varying inputs. We defined the inherently “unexpected” input as a randomly applied set of impulses. The sum of the sinusoid and impulse inputs was then applied to demonstrate a case where different dynamical sources can be separated.

### Decomposition into Optimal Subspaces (DOS) algorithm

We based DOS on a published method that jointly optimizes subspaces separating preparatory and movement activity within motor cortex^19^. We designed DOS to identify two mutually orthogonal subspaces within the neural manifold, one capturing activity that covaried with the neural manifold of another brain region and the other capturing activity that covaried with behavior. Our method looks for eigenvectors spanning subspaces of the neural manifold that maximize the covariance with external covariates (e.g., behavioral covariates). The relevant value to be optimized is defined as:

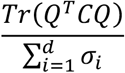

Where:

*Q*: basis vectors of the subspace

*C*: covariance of neural activity and external covariates

*d*: dimensionality of subspace

*σ*_*i*_: variance explained in the neural population by each subspace dimension

The numerator effectively measures the amount of covariance between neural activity and external covariates captured by the subspace. We defined *C* as the covariance between a matrix of *N*-dimensional covariates and *N*-dimensional neural population activity, which we can express as *Y*^*T*^*X*. This formulation ignores a global scaling term for covariance since we are deriving unit basis vectors.

Using this metric, a cost function can be defined that optimizes for any arbitrary number of subspaces, with any value of *d* for each subspace, so long as the sum of their dimensionality is less than or equal to *N*. In our analysis, we defined two subspaces. For the behavioral subspace, *Y* was the activity within the 20-dimensional behavioral manifold identified from the behavioral covariates described above. For the feedback subspace, *Y* was the activity within the 20-dimensional somatosensory cortical manifold. These were referenced against *X*, the 20-dimensional activity within the motor cortical manifold.

We defined the cost function using the above metric for the two subspaces:

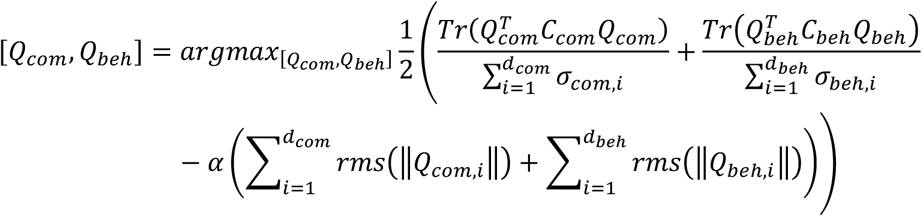

Subject to the constraints:

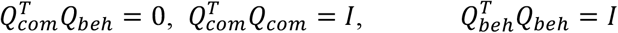

Where:

*Q*_*com*_: basis of feedback subspace

*Q*_*beh*_: basis of behavioral subspace

*X*: local neural population activity (e.g. motor cortex)

*Y*_*neur*_: neural population activity of other brain region (e.g. somatosensory cortex)

*Y*_*kin*_: kinematic and behavioral covariates

*d*_*com*_: dimensionality of the feedback subspace

*d*_*beh*_: dimensionality of the behavioral subspace

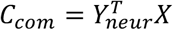: covariance of local neural population activity with other brain region

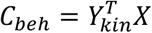: covariance of local neural population activity with behavioral covariates

The first two terms of the cost function maximize the ability of the feedback subspace to explain the covariance of the neural population with the other brain region, as well as the ability of the behavioral subspace to explain the covariance of the neural population with the kinematic signals. The last term penalizes sparsity in the subspace weights. This term is maximized when the weight terms are evenly distributed across the dimensions. In practice, our results on the neural datasets were qualitatively similar without this sparsity constraint, however from our simulation experiments described below, this term was important to avoid edge cases. For example, if a high-variance principal component can explain a lot of behavioral covariance, the algorithm may settle into local minima by attributing a high weight to one dimension. Instead, we want to bias towards finding distributed representations across dimensions. Typically, such sparsity constraints are achieved using the L1-norm. However, since the algorithm optimizes for a basis set, the L1-norm of Q is always 1 regardless of the weights. To capture the same intuition of penalizing sparse weightings to a basis set, we opted to use the root-mean-square of the weights. The *α* term helps to balance the contribution of the different terms.

The cost function used here can be viewed as analogous to that of Reduced Rank Regression, which specifically finds dimensions within the full population space that covary with other signals^13^. It is important to note that there is no constraint in the cost function to explain neural variance, only to explain the *co*variance with external signals. In theory, one of the subspaces could actually explain very little neural variance. The optimization was performed using a Matlab-based manifold optimization framework^51^. Thus, our algorithm can jointly optimize multiple orthogonal subspaces within the same neural population, each subspace designed to explain the covariation of the full population with external signals. A key assumption in the above algorithm is that the dimensionality is matched across all sets of signals, i.e., that *C*_*beh*_ and *C*_*com*_ are square and of identical size. We achieved this by performing principal component analysis on the population activity and kinematic signals before running this algorithm and selecting the same number of dimensions for all sets of signals. We used DOS to identify motor cortical feedback and behavioral subspaces from neural activity and behavior covariates from normal trials only. To analyze the error trials, we projected the neural activity from the error trials onto these subspaces identified from normal trials. This ensured that the error analysis results were not biased by the optimization algorithm.

We quantified the amount of variance captured by each of the behavioral and feedback subspaces using the eigenvalues of the covariance matrix between the subspaces and the activity in the motor cortical manifold.

### Simulation to validate the DOS algorithm

We simulated population of spiking motor and somatosensory cortical neurons where the ground truth latent dynamics are known to confirm that the above decomposition algorithm can correctly reconstruct the latent dynamics in the intended subspaces. We generated two sets of latent dynamics from a one-dimensional random walk. For comparison with the neural and behavioral data from the paper, we refer to these components as feedback and behavioral, although as random processes they do not have any such structure inherently built in. We then smoothed these random dynamics with a Gaussian kernel of 50ms width to obtain smoothness that is comparable to that of the neural data. We developed DOS with intention to identify specific patterns of covariance, even in the face of confounding correlated signals. To test the accuracy of the algorithm, we added two confounds. First, both the motor and somatosensory cortical populations received correlated dynamics (modeled as sinusoids) to simulate the effect of common input to both regions (which were referred to as null dynamics). These are important to induce potentially spurious correlations in the dataset which are not directly related to the variables of interest. Second, the somatosensory cortical population received the precise input-driven dynamics as well as a time-shifted version of the deterministic dynamics meant to simulate the real-world delays between intended motor output (here assumed to be 50ms) and the corresponding sensory feedback (assumed to be another 50ms).

The motor and somatosensory cortical population activity was generated as a Poisson process whose input was randomly weighted combinations of the null, input-driven, and deterministic dynamics. We then processed the simulated spiking activity with the same methods as the real neural data, including normalization, smoothing, and dimensionality-reduction through PCA. We identified the subspaces within the simulated motor cortical data using the covariance between motor and somatosensory cortical PCs, as well as the covariance between motor cortex and the known deterministic dynamics. We quantified the performance by computing the variance explained (R^2^) between the ground truth dynamics and the extracted subspace dynamics. We repeated this random simulation 1000 times to generate the performance distributions. Note that by design DOS cannot separate highly correlated inputs. Thus, on each candidate random simulation we computed the Pearson’s correlation between the two sets of latent dynamics and rejected simulations with values greater than 0.25. On each simulation, we compared the performance of DOS to a null distribution found by taking the max correlation across all individual motor cortical PCs. This null distribution served to confirm that the reconstructed dynamics did not trivially arise from covariance across neurons found by PCA, but instead required a targeted approach like DOS.

### Behavioral task

The monkeys were seated in a custom primate chair that enabled unconstrained interaction with a workspace in front of their body. The experimental platform and task has been previously described in detail in ^10^. In brief, a small, spherical, custom-molded, silicon object was affixed to the end of a seven-degree-of-freedom robotic arm (Intelligent Industrial Work Assistant, IIWA – KUKA, Augsburg, Germany). Custom control software enabled positioning of the robot in space. For Mk-Jo, only one position was provided, roughly centered in front of the body. For Mk-Cs, Mk-Br, and Mk-Ol, three positions were used: left, middle, and right. All positions were in the same coronal plane. The monkeys were trained to freely reach for the object, grasp it using only their left hand, and then pull it towards the body. In each trial, the robot brought the object to one of three randomly-selected positions (except Mk-Jo, where only one position was used). While the object was displaced from its starting position, the robot provided a force towards the starting position that was proportional to the horizontal displacement. Therefore, the monkeys had to resist this force to pull the object and, if they released it, the object quickly returned to its starting position. The system recorded the interaction forces produced by the robot.

Each trial was deemed successful after the position of the object passed a pre-determined distance threshold. Pulling movements that failed to reach this threshold were excluded. Upon success, Mk-Cs and Mk-Jo manually received a food reward, while Mk-Br and Mk-Ol received an automated liquid reward through a sipper tube. Error trials were manually identified by assessing the videos and kinematic trajectories. Most of the errors occurred when the monkey missed the robot in the initial reach or when their hand slipped off the robot while pulling. Typically, this resulted in errors preceding the final successful grasp, though for Mk-Br most errors occurred after grasping the object to adjust the posture of the hand to enable the pulling movement. After final curation, we analyzed the following number of trials: Mk-Cs: 100 normal, 22 error; Mk-Br: 158 normal, 66 error; Mk-Ol: 234 normal, 88 error; Mk-Jo: 27 normal, 6 error. While performing the task, the monkeys were affixed with up to 15 reflective markers to track kinematics of the joints of the arm and hand (Vicon Motion Systems, Oxford, UK).

### Surgical procedures for cortical implants

All surgical procedures were performed using aseptic technique under general anaesthesia (induction: 0.1 mg/kg midazolam and 10 mg/kg ketamine; maintenance: 5 ml/kg/h propofol and 0.2-1.7 ml/kg/h fentanyl IV). A certified neurosurgeon (Dr. Jocelyne Bloch, CHUV, Lausanne, Switzerland) supervised all procedures. Each monkey was implanted with two Utah electrode arrays (Blackrock Microsystems, Salt Lake City, UT, USA). The first array targeted the arm region of motor cortex near the central sulcus (primary motor cortex; 1.5 mm shaft length) and the other targeted somatosensory cortex (Brodmann’s Area 2; 1 mm shaft length). The two arrays were connected to a single pedestal mounted to an implanted titanium that covered a portion of the skull. The mesh was molded to the shape of the skull of each monkey using a plastic skull 3-D printed from an individualized MRI scan. The implants in Mk-Cs and Mk-Br targeted arm areas identified by intraoperative surface stimulation (biphasic pulses, 3mA), while the implants in Mk-Jo targeted the hand areas. All arrays had 64 channels in an 8×8 configuration except the somatosensory cortical array of Mk-Br, which was 32 channels in an 8×4 configuration. Mk-Br and Mk-Ol each additionally received a third array (32-channel, 1mm shaft) anterior to the first array near to ventral premotor cortex. We saw qualitatively similar dynamics in this and the array implanted near the Central Sulcus. We therefore combined the signals recorded from both arrays in each monkey and for simplicity refer to this combined signal as “motor cortex”. In the same surgery, Mk-Br also received bipolar EMG electrodes in the left arm to record the activation of eight muscles of the arm and hand. Detailed surgical and post-operative care procedures have been described previously^52^.

### Neural data acquisition

As the monkeys performed the behavioral task, we recorded infrared video from 12 cameras to track the kinematic markers at 100Hz (Vicon Motion Systems, Oxford, UK) and neural activity at 30kHz using a Cerebus system (Blackrock Microsystems, Salt Lake City, UT, USA). To process the neural data, we bandpass-filtered each channel at 750 to 5000 Hz and set a threshold between -5x and -6.25x the RMS value to extract spike events. We sorted these spikes using Offline Sorter (Plexon, Dallas, TX, USA) to identify putative single neurons. We counted the number of spikes occurring in 10ms bins matched to the 100 Hz kinematic data. We used these binned spike counts in the Generalized Linear Model analysis (see below). For all other analyses, we square-root transformed each spike train to stabilize the variance^53^ and then converted them to an instantaneous firing rate by convolution with a Gaussian kernel with a 75ms standard deviation.

### Calculation of neural and behavioral trajectories

We applied a soft normalization procedure^6^ to all of the smoothed single neuron firing rates using a fixed constant of 5 in the denominator. We chose to normalize to ensure fair comparison between different neural and behavioral manifolds, for example to compensate for the typically lower firing rates of somatosensory cortical neurons^10^. In practice, our results were unchanged with reasonable adjustments to this value. We then used PCA to reduce the dimensionality of each brain region independently and compute a neural manifold^20^. We selected the first 20 dimensions, though in practice our results were not impacted by the exact choice of dimensionality within reasonable ranges (10-30 dimensions; data not shown). We then computed a “behavioral manifold” using a similar procedure. First, we z-scored each of the position, velocity, acceleration, and force signals for the limb and robot. For Mk-Br, we also added the EMG recordings to this behavioral manifold to most accurately estimate the behavioral dynamics. We then applied PCA to this set of signals and selected the first 20 dimensions to correspond to the 20 dimensions of the neural manifold. The match in dimensionality was necessary for the subspace decomposition analysis described below, as well as to compare the trajectory tangling between manifolds.

### Computing trajectory tangling

We employed trajectory tangling to quantify the relative dynamical behavior of the neural and behavioral dynamics. Tangling values should be highest at points in time where the current state could lead to multiple future states. We computed the instantaneous tangling (*T*) at each time *t* of the neural and behavioral trajectories as described previously^9^.

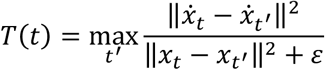

The metric looks over all time points for all trials for each position and is maximal when a similar position in state space (*x*) – which gives a small denominator value – corresponds to different state space velocities 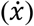 – which gives a large value in the numerator. Such states would be expected to occur when the system is driven by unexpected inputs – causing a discrepancy between the past velocity and the future evolution – or when the system is chaotic or unstable and linear dynamical systems assumptions break down. We do not consider the latter case to be likely in practice in our motor cortical data given the widely demonstrated robustness of cortical signals. Thus, we largely interpret increases in tangling as an increase in the influence of unexpected inputs

The value of *ε* was chosen to be very small (10^−6^) and served only to keep the value from becoming undefined if the denominator was otherwise equal to zero. However, *ε* could also be scaled based on the signal magnitude as in Russo et al.^9^, though we observed qualitatively equivalent results when we took this approach. We computed the tangling at firing rate estimates in 10ms bins. Note that we computed the tangling independently for all trials for each position, though we obtained similar results computing the tangling across all trials together. In previous work, tangling has been applied to trial-averaged data. Thus, noise and outlier datapoints were much less likely. We modified the approach slightly to take the 99.99^th^ percentile across all time points in each position instead of the max in order to ignore a very small number of outliers (e.g. artifacts) in our single trial data.

Tangling values are sensitive to the dimensionality of the state space and, therefore, can be directly compared only between spaces with matched dimensionality. This effect is explored in depth in **Fig. S7**, where we demonstrate that in our datasets tangling monotonically decreases as assumed dimensionality increases. To facilitate easy comparison across signals or conditions, we ensured that dimensionalities were matched (20 dimensions for manifolds, 5 dimensions for subspaces). To study the time course of tangling changes, we linearly interpolated to time-warp each trial to be the same length with a basis of 200 samples, preserving the proportion of time spent on average in the reaching and pulling phases. We then averaged across normal or error trials (e.g., **Fig. 3d**) To compute an aggregate single-trial tangling value, we computed the root-mean-square tangling through the trial (e.g., **Fig. 3b**) or throughout the specific period of behavioral error correction (e.g., **Fig. 3e**).

### Modeling deterministic dynamics without inputs to provide reference values for tangling

We designed data-constrained RNNs to understand the components of motor and somatosensory cortical recordings that can be explained by purely deterministic dynamics. Unlike the previous RNN models, which were randomly connected, chaotic networks, these data-constrained RNNs were specifically trained to generate stable, autonomous (i.e. not input-driven), deterministic dynamics in a manner consistent with recorded experimental data. The models were trained according to the methods in Ref. ^32^. In brief, we instantiated a randomly connected matrix from a Gaussian distribution with a number of units equal to the number of recorded neurons to give a 1:1 correspondence. The weights were initially scaled by *g*=1.3, where *g* is a parameter related to the chaos in the network. The models were then trained iteratively through a variant of Recursive Least Squares^32,54^ with a cost function that minimized the error between the activity of each RNN unit and its corresponding recorded neuron. Once trained, the model RNN autonomously and deterministically generated dynamics consistent with the experimental data. Importantly for the purposes of this paper, the model RNN was only able to capture the aspects of the motor and somatosensory cortical population activity that can be well-described using a single, autonomous dynamical system. Any periods of strong input-driven dynamics will not be well captured by the model. Since the RNNs were fit to each monkey individually and were tied to the specific features and dimensionality of the real data, they provided a meaningful and interpretable baseline value for what values to expect from the tangling analysis when only deterministic dynamics are present in the system. We thus computed the tangling on individual trials using the same methodology and processing as in the real motor cortical data (**Fig. 3e**). This includes, for the case of the motor cortical population activity, the gaussian smoothing of the RNN unit responses to ensure parity with the neural recordings. We also performed an identical subspace decomposition and re-computed the tangling in each of these subspaces to use as a reference for the real subspaces (**Fig. 4d,e**).

### Spinal cord stimulation experiments to validate subspace decomposition

Portions of the spinal stimulation datasets were previously published in Ref. ^33^ In brief, two monkeys (Mk-Br and Mk-Yg) received the cortical implants described above. In a separate surgery following recovery, the monkeys were chronically implanted with flexible e-dura spinal stimulation implants in the epidural space covering spinal segments C6 to T1, that routed to an additional connector attached to the skull. A focal lesion targeting the descending corticospinal tracts was then performed at the C5/C6 level above the stimulation implant. After recovery, the subjects exhibited pronounced deficits in distal arm and hand functions but largely retained control of the proximal muscles of the arm. We recorded from the implanted cortical arrays while they attempted to perform the reach, grasp, and manipulate task described above, regardless of whether they were successful on the trials. This created a dataset of attempted residual movements with which to define the feedback and behavioral subspaces, as in the previous experiments. We then began to stimulate below the lesion using the implanted spinal arrays. We stimulated with irregular timing as the monkeys attempted to complete the task, allowing some pulse to overlap with periods of rest where no movements occurred.

We applied DOS to identify putative behavioral and feedback subspaces using only the attempted reaching movements where no stimulation was applied. We then selected the stimulation periods where we could verify that the arm has near the rest position and had zero velocity throughout the duration of the stimulation. The stimulation frequency for the trials considered in the following analyses (N=68 for Mk-Br and N=37 for Mk-Yg) ranged from 20-100Hz and lasted for 500ms. The stimulation pulses were applied at various spinal segments ranging from C6 to T1. For the purposes of this analysis, we pooled together stimulation applied to all sites with the logic that all stimulation attempts were below the lesion and would recruit overlapping sets of ascending dorsal column fibers.

We aimed to quantify the effect of the stimulation on neural activity in the two subspaces. We aligned the neural activity on the onset of each stimulation attempt and projected the motor cortical population activity into each subspace. Note that, unlike the previous scenarios, it was challenging to apply the trajectory tangling metric in this experiment since the monkey was at rest during the stimulation. Trajectory tangling during rest is by its nature ill-defined, and any intervention that causes the trajectory to deviate from rest will necessarily cause a transient increase in tangling regardless of the underlying dynamical features of the system. Instead, we computed the change in total variance within each subspace over time relative to the value at the onset of stimulation (**Fig. 5e**). We then assessed the temporal evolution of responses in each subspace by comparing the canonical correlation (canoncorr in Matlab, The Mathworks, Inc.) between the trajectories at different temporal shifts, averaged across the first three aligned dimensions (**Fig. 5f**).

## SUPPLEMENTARY FIGURES

**Figure S1.**
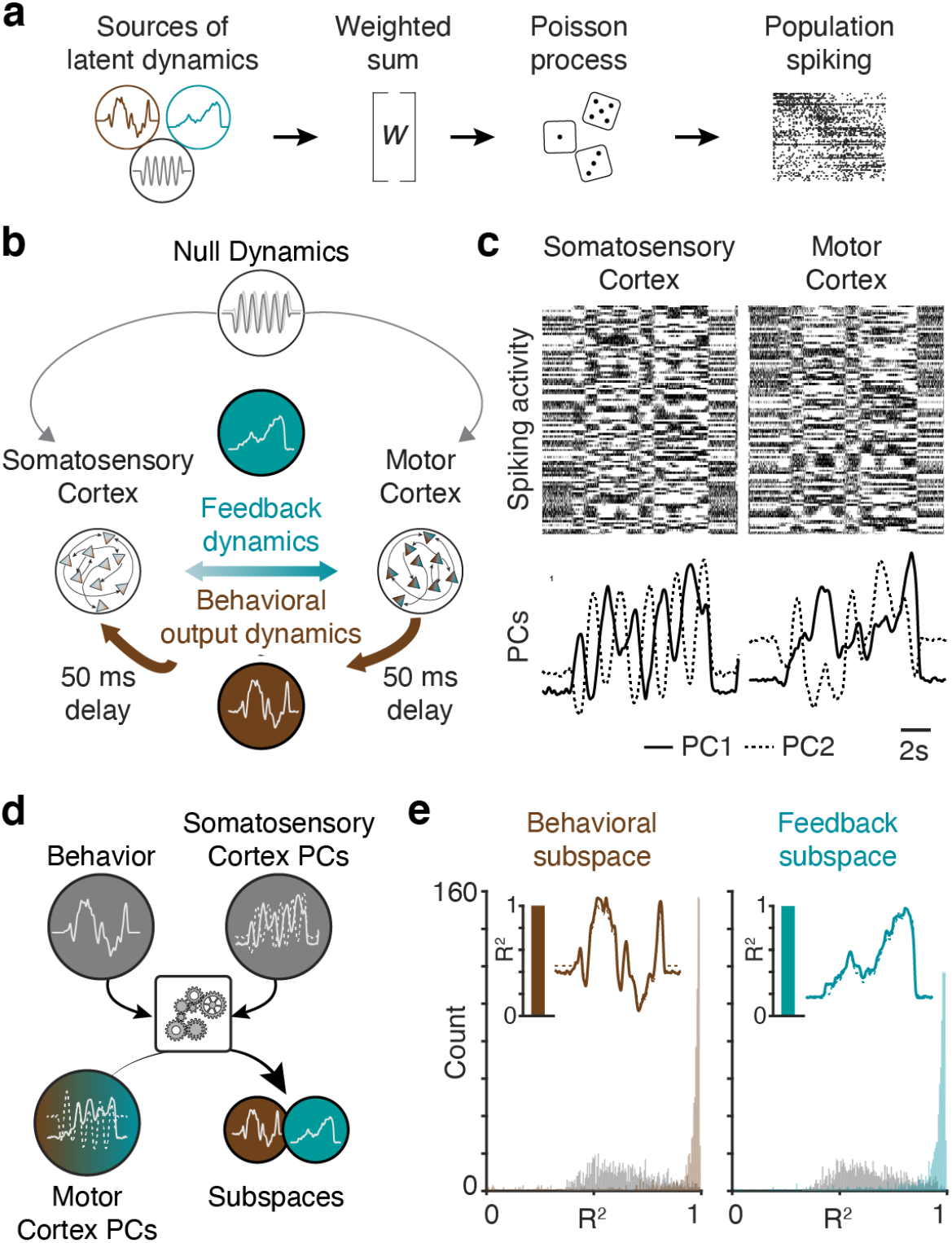
The Decomposition into Optimal Subspaces (DOS) algorithm accurately isolates subspaces from simulated neural activity. **(a)** We modeled neural activity of 100 motor and somatosensory cortical neurons whose activity was generated by randomly weighted combinations of sources of latent dynamics. **(b)** We defined three types of latent dynamics from a random walk process. While the outputs were random, we gave these random dynamics names to correspond to the dynamics studied in the neural and behavioral datasets.: (i) so-called “feedback” dynamics shared between both regions, (ii) “behavioral” dynamics that were also correlated between regions but delayed in somatosensory cortex to account for sensory input-driven delays, and (iii) oscillatory null dynamics intended to confound the algorithm. **(c)** We applied PCA on the spiking activity of the simulated neurons (top) to reduce the dimensionality of the dataset (bottom shows the two leading Principal Components, PCs). Each PC reflects a mix of the three sources of dynamics, thus PCA alone is not sufficient to separate the dynamics. **(d)** The DOS algorithm aimed to isolate the feedback and behavioral dynamics into orthogonal subspaces by referencing the motor cortical activity against the behavioral dynamics (assumed to be measurable as in the experimental data), and the somatosensory cortical manifold. **(e)** For the example simulation shown in Panels a-c, the algorithm achieved near-perfect reconstruction (R^2^ > 0.95) of the behavioral and feedback dynamics in the two orthogonal subspaces (insets). We performed 1000 such simulations and the algorithm consistently performed well (median R^2^: 0.97), significantly better than the max across all PCs in gray (p < 0.01, Mann-Whitney signed rank test). (Left) The histogram shows the distribution of the R^2^ values calculated between one of the PCs (grey) or the behavioral subspace (brown) and the behavioral dynamics. The inset plot shows the behavioral dynamics (brown line) and the behavioral subspace for one simulation. The inset bar plot shows the R^2^ value for that example. (Right) The histogram shows the distribution of the R^2^ values calculated between one of the PCs (grey) or the feedback subspace (brown) and the feedback dynamics. The inset plot shows the feedback dynamics (brown line) and the feedback subspace for one simulation. The inset bar plot shows the R^2^ value for that example.

**Figure S2.**
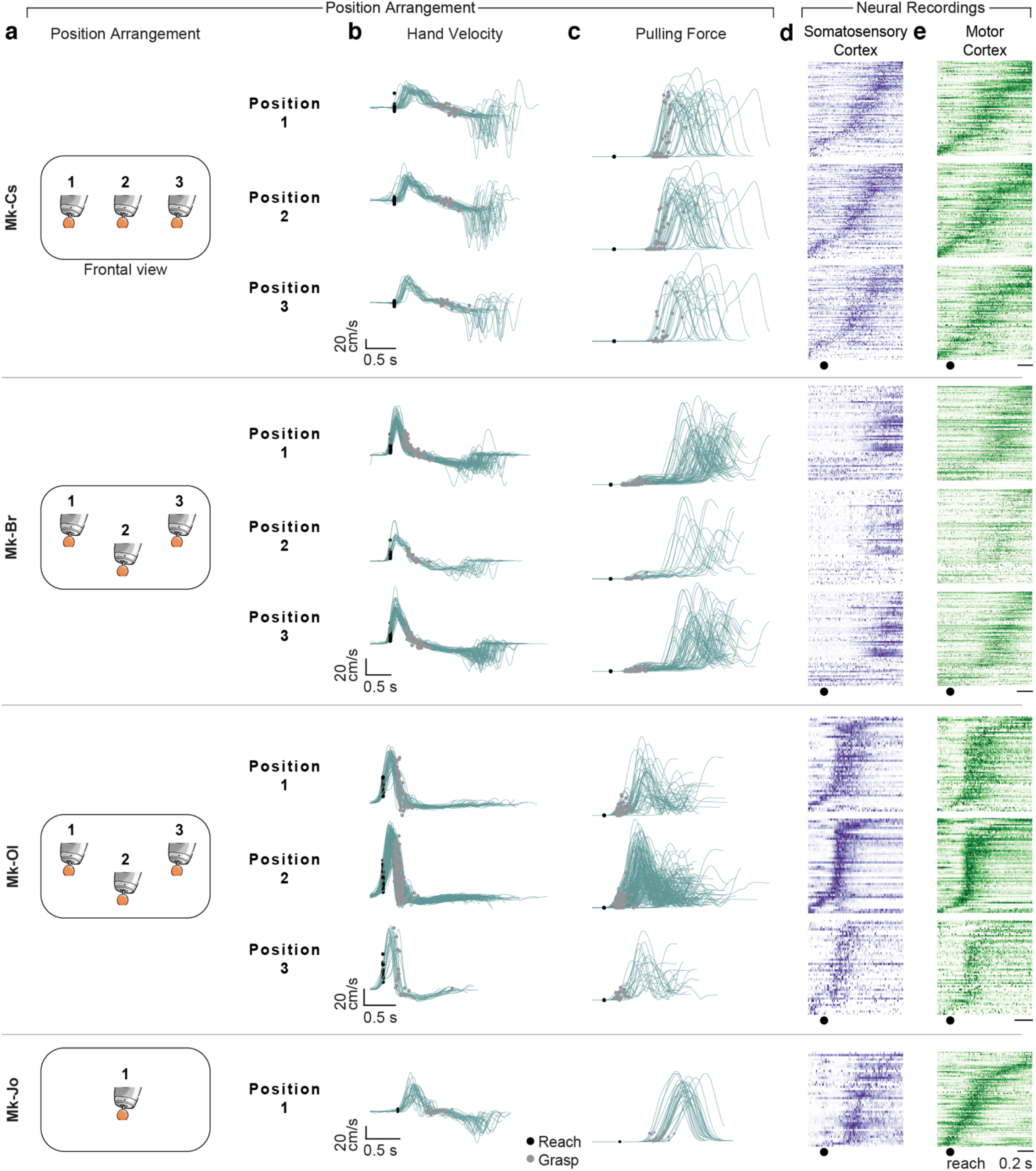
Detailed description of behavioral and neurophysiological dataset. **(a)** Arrangement of object positions in the coronal plane for each monkey. Mk-Jo reached to only one position in the center of the workspace, while the other three monkeys reached to three positions. **(b)** Plots show hand velocity traces for each position for all monkeys. Trajectories are duplicated from Fig. 2c, but separated by position. Black circles denote the velocity in the sagittal plane at the time of reach onset marked from the video recordings, and gray circles denote velocity at the time of object grasp. **(c)** Plots show the magnitude of pulling force registered by the force transducer in the robot for each position. **(d)** Peri-event histograms for somatosensory cortex spiking activity (spikes counted in 10ms bins aligned on reach onset). Each row represents the sum across trials of one neuron. **(e)** Data presented as in Panel d, but for motor cortical neurons.

**Figure S3.**
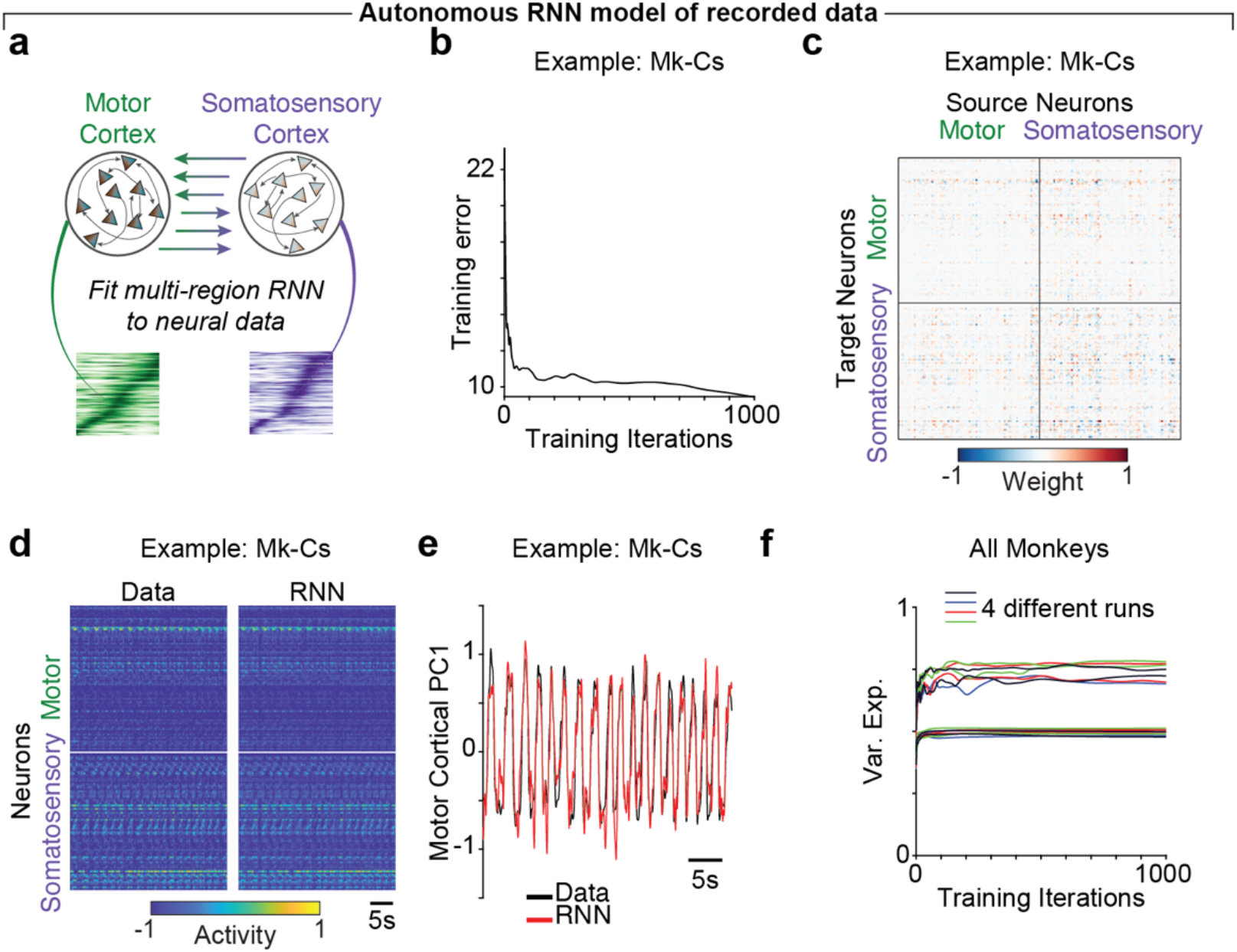
Data-driven autonomous RNN models of motor cortical activity provide context on tangling values. **(a)** We designed RNN models that were directly constrained by the time-series neural recordings from motor and somatosensory cortex for each monkey. **(b)** Example training error over 1000 iterations for Mk-Cs. **(c)** Heatmap of weights inferred by the RNN for the interactions between each neuron for Mk-Cs. Values were normalized across rows to aid visualization. **(d)** Heatmap comparing 22 trials for Mk-Cs for the recorded data (left) and the RNN model (right). **(e)** Projection of motor cortical population activity onto the first PC for the recorded data (black) and the RNN model (red) for Mk-Cs. **(f)** Variance explained in the neural population by the RNN fits for different random initializations across four independent training runs for each of the four monkeys (16 lines). The autonomous RNNs saturated at approximately 50% to 80% explained variance, suggesting that deterministic dynamics cannot capture all of the dynamics present in the motor and somatosensory cortical activity.

**Figure S4.**
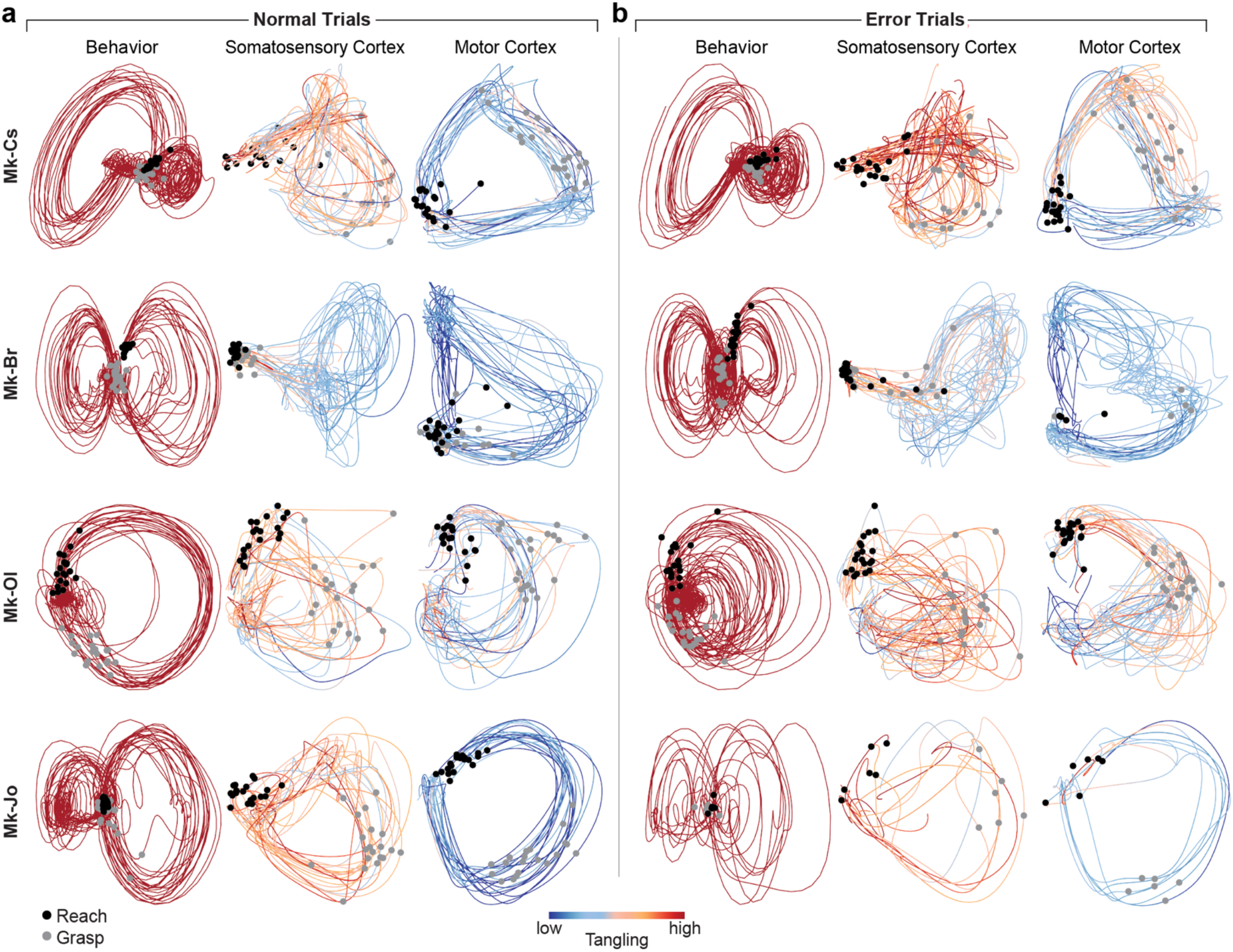
Comparison of the trajectories and tangling between behavioral, somatosensory cortical and motor cortical manifolds during normal and error trials. **(a)** Trajectories of the behavioral manifold are substantially more tangled than the somatosensory cortical manifold trajectories, which are substantially more tangled than the motor cortical manifold trajectories on normal trials. Plots show trajectories on 20 normal trials for behavioral (left), somatosensory cortical (middle), and motor cortical manifolds (right) shown in the two leading PCs for the four monkeys (four rows). Colors indicate tangling value at each time point on a logarithmic scale, where blue and red correspond to the minimum and maximum neural tangling across all time points for the motor cortical and somatosensory cortical manifolds of all normal and error trials for each monkey, respectively. The tangling values for the behavioral manifold are allowed to saturate. Data are reproduced from Figure 3a. **(b)** Tangling in both somatosensory and motor cortical manifolds increases during the error trials. Plots show trajectories for up to 20 error trials for behavioral (left), somatosensory cortical (middle), and motor cortical manifolds (right) for the four monkeys. Color scale is identical for each monkey to the scale in Panel a.

**Figure S5.**
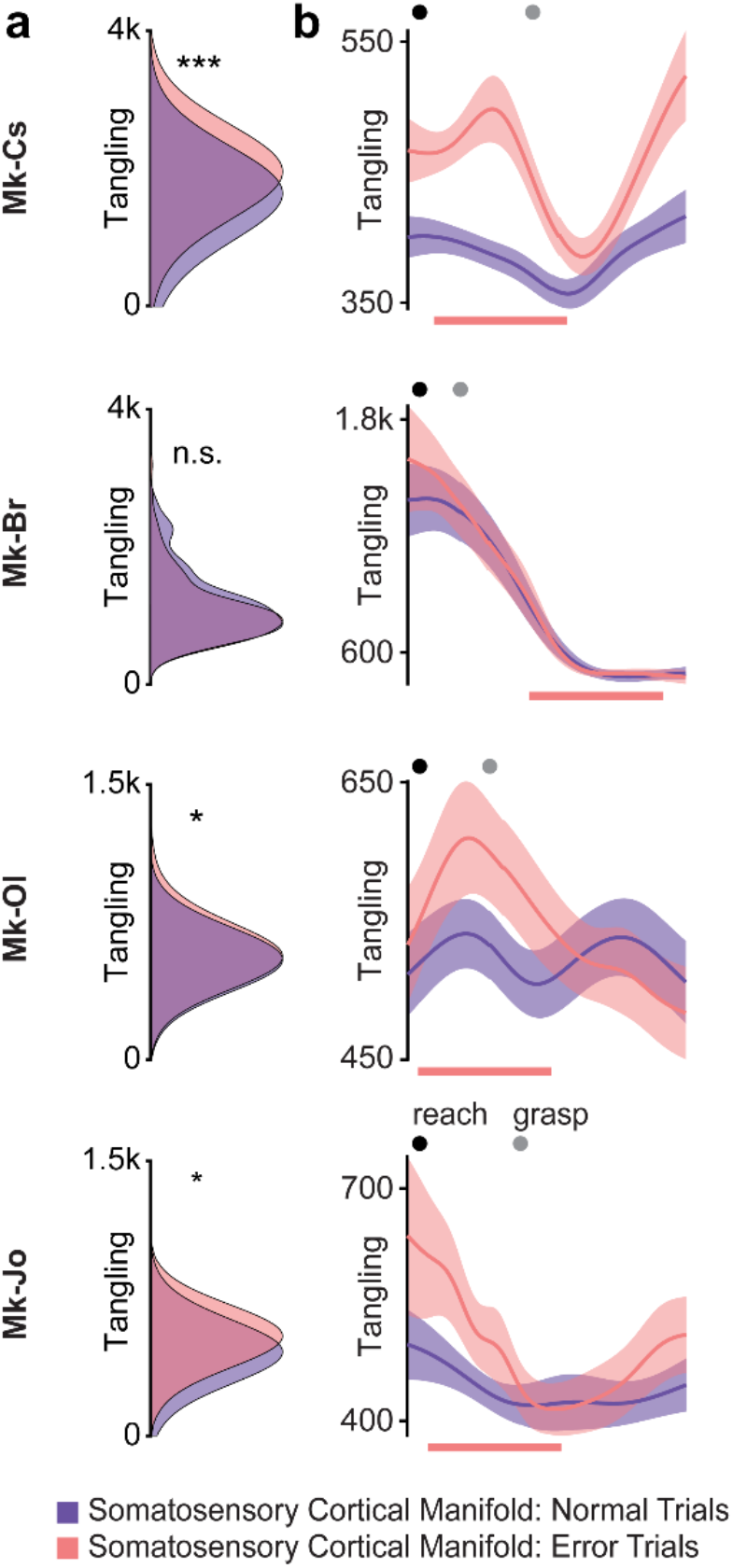
Errors lead to a slight increase in the tangling of somatosensory cortical manifold. **(a)** Plots show single-trial tangling distributions in the somatosensory cortical manifold for all normal (purple) and error (pink) trials. All monkeys except Mk-Br showed a significant increase in tangling on error trials, though Mk-Br did occasionally show small, transient increases in tangling (see Fig. S4b). *: p < 0.05, **: p < 0.01, ***: p < 0.001; ranksum. **(b)** Somatosensory cortical tangling increases surrounding the error correction in all but one monkey. Tangling in the somatosensory cortical population averaged across all normal (purple) and error (pink) trials after aligning on reach onset and time-warping to match the length across all trials. Black and grey circles indicate the reach onset and object grasp, respectively. Pink bars indicate approximate windows of error correction for each monkey. Error bars: mean ± s.e.m.

**Figure S6.**
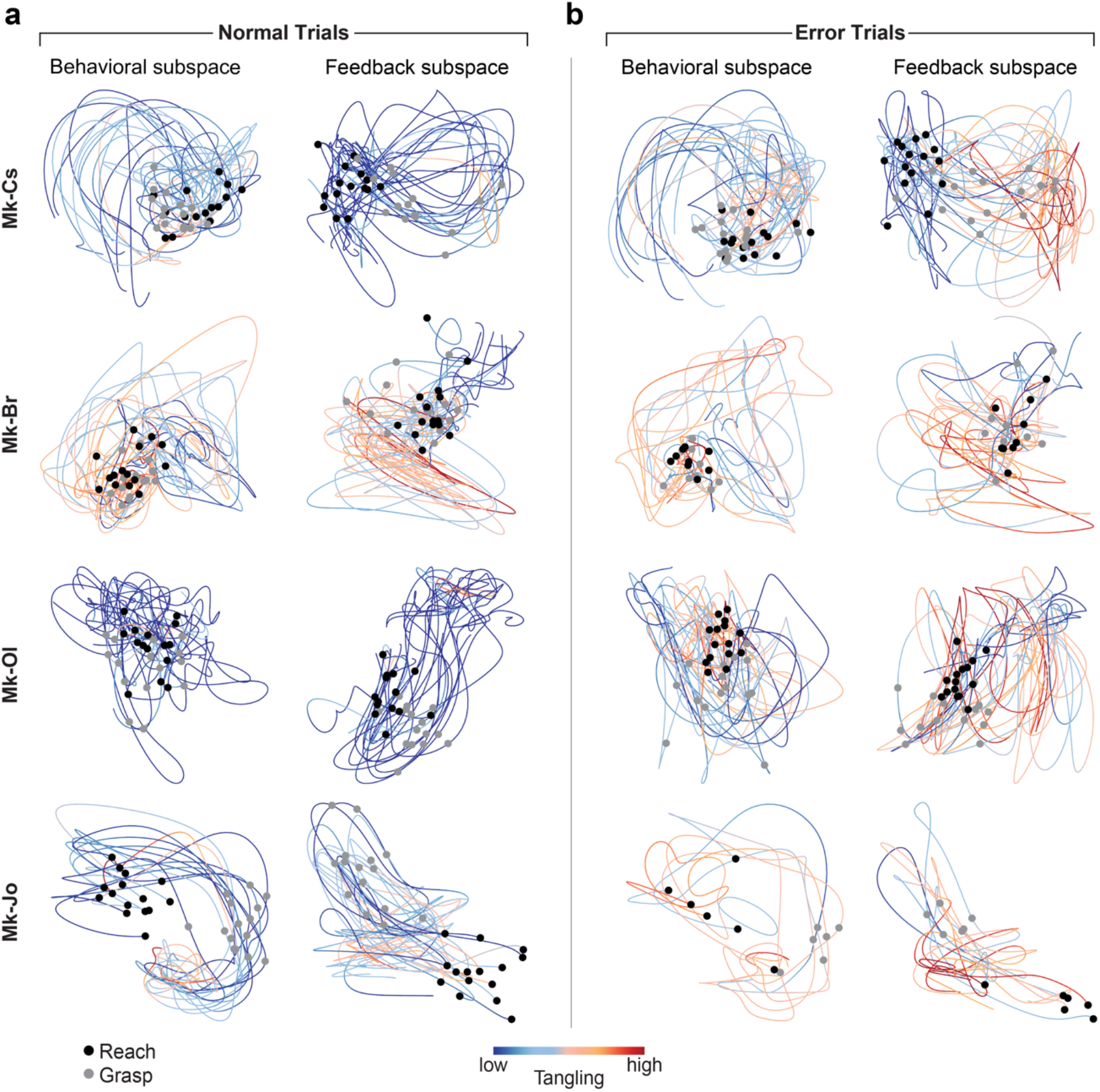
Tangling of feedback subspace trajectories increases substantially during errors compared to normal trials, while the tangling of behavioral subspace trajectories does not change. **(a)** Neural trajectories within the behavioral (left) and feedback (right) subspaces for 15 normal trials for the four monkeys. Data plotted as in Figure 6a; Mk-Cs trajectories reproduced from that figure. **(b)** Neural trajectories in the behavioral (left) and feedback (right) subspaces for up to 15 error trials for the four monkeys. Color scale is consistent with Panel a.

**Figure S7.**
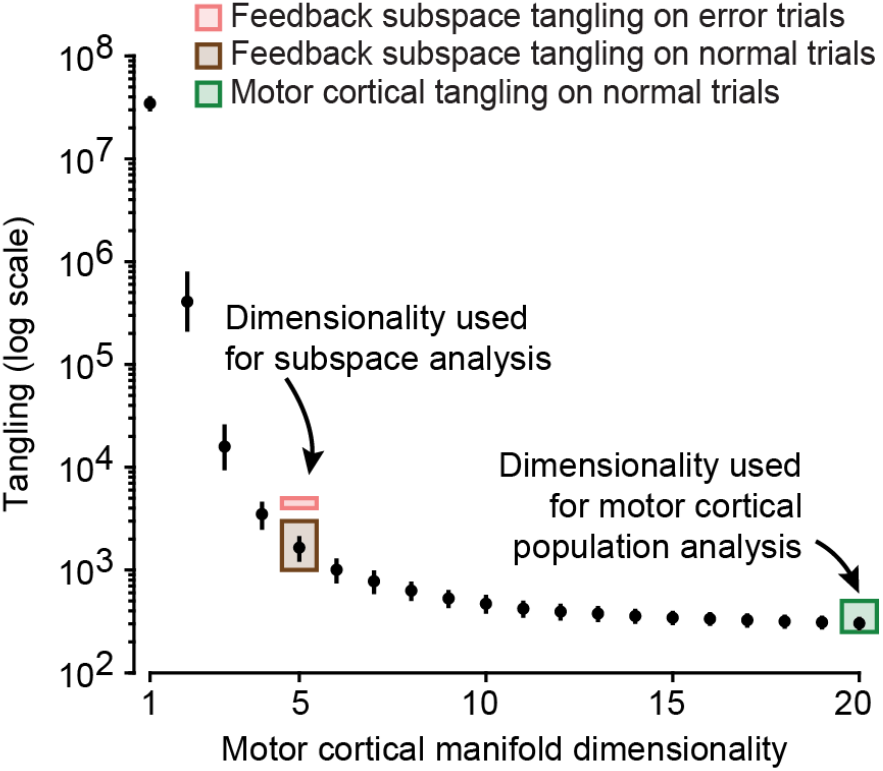
Tangling is dependent on the dimensionality of the manifold. We repeated the tangling analysis in the motor cortical manifold at a range of dimensionalities. As expected, given that tangling is a relative metric, the values monotonically decreased as dimensionality increased. However, the choice of dimensionality alone could not explain the increased tangling we observed on error trials in the feedback subspace on error trials.

**Figure S8.**
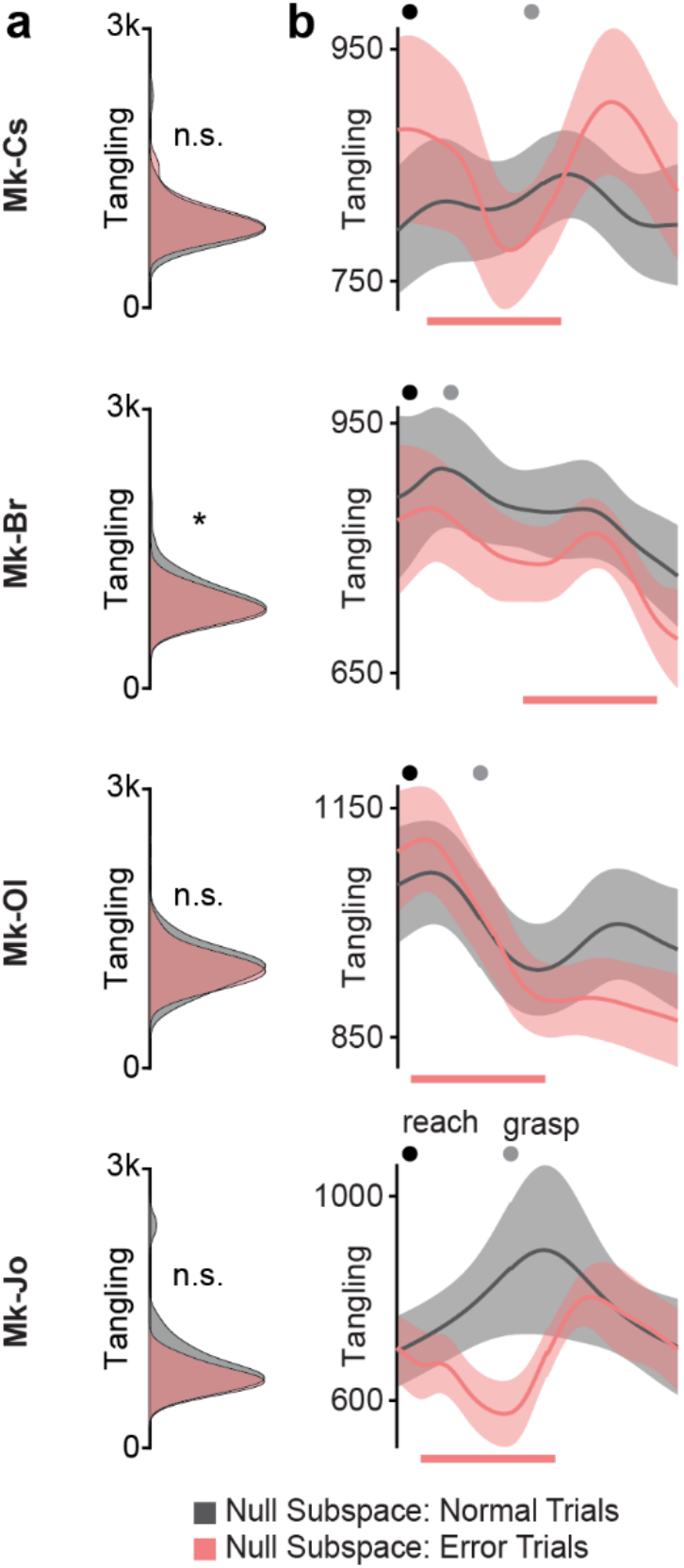
Errors lead to no increase in the tangling of the null subspace of the motor cortical manifold. We used DOS algorithm to identify five-dimensional feedback and behavioral manifolds. The remaining ten dimensions of the motor cortical manifold activity spanned the null space. **(a)** Single-trial tangling distributions in the null subspace for all normal (gray) and error (pink) trials. No significant difference was observed for Mk-Cs, Mk-Ol, or Mk-Jo, though Mk-Br showed a small yet significant decrease on the error trials. n.s.: not significant, p ≥ 0.05; *: p < 0.05; ranksum. **(b)** Plots show tangling in the null subspace (see Methods) averaged across all normal (gray) and error (pink) trials after aligning on reach onset and time-warping to match the length across all trials. Black and grey circles indicate the average time of reach onset and object grasp, respectively. Pink bars indicate approximate windows of error correction or each monkey. Data plotted as mean ± s.e.m.

